# Realizing Mechanical Frustration at the Nanoscale Using DNA Origami

**DOI:** 10.1101/2024.06.26.600849

**Authors:** Anirudh S. Madhvacharyula, Ruixin Li, Alexander A. Swett, Yancheng Du, Friedrich C. Simmel, Jong Hyun Choi

**Affiliations:** School of Mechanical Engineering, West Lafayette, Indiana 47907, USA; School of Materials Engineering, Purdue University, West Lafayette, Indiana 47907, USA; Physics of Synthetic Biological Systems, School of Natural Sciences, Technical University of Munich, Garching 85748, Germany

**Keywords:** DNA origami, self-assembly, metamaterials, geometric frustration, mechanical design, mechanics, free energy landscape

## Abstract

Structural designs inspired by physical and biological systems have been previously utilized to develop advanced mechanical metamaterials. These are based on the clever geometric arrangement of their building blocks, resulting in enhanced mechanical properties such as shape morphing and auxetic behavior. Until now, the benefits from such designs have yet to be leveraged at the nanoscale. Here, we use the DNA origami method to realize a nanoscale metastructure exhibiting mechanical frustration, the mechanical counterpart of the well-known phenomenon of magnetic frustration. We show that this DNA metastructure can be precisely controlled to adopt either frustrated or non-frustrated mechanical states, each characterized by a distinct free energy profile. Switching among the states is achieved by engineering reconfigurable struts into the structure. Actuation of the struts causes a global deformation of the metastructures. In the non-frustrated state, strain can be distributed homogeneously throughout the structure, while in the frustrated state, strain is concentrated at a specific location. Molecular dynamics simulations reconcile the contrasting behaviors of the two modes and provide detailed insights into the mechanics. Our work demonstrates how combining programmable DNA self-assembly with mechanical design principles can overcome engineering limitations encountered at the macroscale, enabling the development of dynamic, deformable nanostructures with tunable responses. These may lay the foundation for mechanical energy storage elements, nanomechanical computation, and allosteric mechanisms in DNA-based nanomachinery.

## INTRODUCTION

Mechanical metamaterials are designed by arranging mechanically coupled, cellular building blocks to achieve nonlinear stress responses^1^, shape morphing^2^, or auxetic deformation^3^. The geometric configuration of the building blocks is key in metastructure design since it gives rise to the observed behaviors. Architected metamaterials cater to a wide variety of applications in aerodynamics^4, 5^, biomedicine^6^, photonics^7^, and acoustics^8^. Among various design strategies, mechanical frustration is a unique approach in the development of mechanical metamaterials, whose behavior is analogous to magnetic frustration. In magnetic frustration, electronic spins cannot interact cooperatively due to geometric constraints imposed by their arrangement in a lattice, resulting in magnetic states with high ground-state degeneracy.^9^ A paradigmatic form of magnetic frustration is exhibited by the Ising model of spins with antiferromagnetic coupling arranged on a triangular or Kagome lattice.^10^ By drawing an analogy between mechanical deformation and electronic spin, similar models can be leveraged to design frustrated metamaterials.^11^ When deformations in neighboring cells of the metamaterial can interact cooperatively, adaptable structures are realized; in contrast, conflicting interactions lead to an inadaptable state where strains are localized. Due to the vast combinatorial number of possible arrangements of the mechanical building blocks, this approach allows for an extremely large design space for structures exhibiting mechanical frustration behaviors. Previous studies following this method used 3D-printed lattices deforming under external forces,^12, 13^ but these are only viable at the meso-to macroscale (i.e., micrometer-sized or larger structures where classical elasticity theories hold). Frustrated metamaterials have never been realized at the nanoscale, because of difficulties in precise assembly and structural accuracy. A design approach that enables mechanical frustration at the nanometer level can open avenues to develop architected nanomaterials with tunable responses, large-scale deformation, and ‘action-at-a-distance’ capabilities.

DNA origami is a bottom-up approach for constructing arbitrary nanostructures based on sequence complementarity^14, 15^. Given the excellent programmability and structural predictability, DNA self-assembly has demonstrated both static and dynamic nanostructures including 2D lattices^16, 17^, 3D polyhedra^18^, reconfigurable switches^19^, kinematic mechanisms^20, 21^, and nanomachines powered by electric fields and chemical potentials^22-24^. Most deformable DNA constructs fall under the umbrella of adaptability where their components interact in synchrony to exhibit overall deformations or even negative Poisson’s ratios^25-27^. Recent reports used computational mechanics to understand relevant properties and thermodynamic models to elucidate free energy landscapes^28, 29^.

Here, we integrate DNA origami with computational design principles to develop nanoscale frustrated metastructures. We design a Kagome lattice using DNA wireframes and calculate the free energies via molecular dynamics (MD) simulations to demonstrate adaptable and inadaptable structures. DNA is used as rigid edges in the wireframe as well as deformable struts. This allows us to dynamically transform between adaptable and inadaptable states, unlike macroscale metamaterials which are normally set by design. Coarse-grained MD models elucidate contrasting behaviors with distinct strain/stress distributions and show how to design metastructures with multistability. We realize the structural design in experiment by using two-step DNA reactions and demonstrate the frustration with buckling edges that can be recovered reversibly.

## RESULTS

### Designing geometric frustration via MD simulation

Drawing inspiration from the Kagome lattice in spin-frustrated systems, we designed a six-cell hexagonal lattice as a model system (Fig. 1a-b) for engineering mechanically frustrated DNA metastructures. Our DNA lattice is composed of honeycomb-patterned, two-helix bundle (2HB) edges with a designed length of ∼28 nm each (see SI for design details, Figs. S2 and S3). To demonstrate both frustrated and non-frustrated states in a single lattice, we embedded additional adjustable struts^30^ shown as colored edges in Fig. 1c which served as actuators in experiment. This structure switches between adaptable and inadaptable modes in response to external loadings applied to the green and blue edges, respectively (Fig. 1d-e). In theory, mapping the deformation of the constituent cells predicts that the adaptable mode will deform freely, distributing the strain throughout the structure, whereas the inadaptable state will be frustrated and localize the strain with buckling edges (Fig. S1). While this is an effective process for designing macroscale metamaterials, nanoscale constructs such as our DNA lattices require additional consideration. This is due to the interplay between thermal fluctuations and the intrinsic mechanical properties of DNA, leading to more complex behaviors. MD simulation is an excellent approach to elucidate this interplay. Coarse-grained models have proven to be successful in explaining structural behaviors of DNA nanostructures^31, 32^. We performed coarse-grained MD simulations using the oxDNA platform^33, 34^. Figure 1f-g shows the equilibrium conformations of the DNA metastructure in the adaptable and inadaptable states.

**Figure 1.**
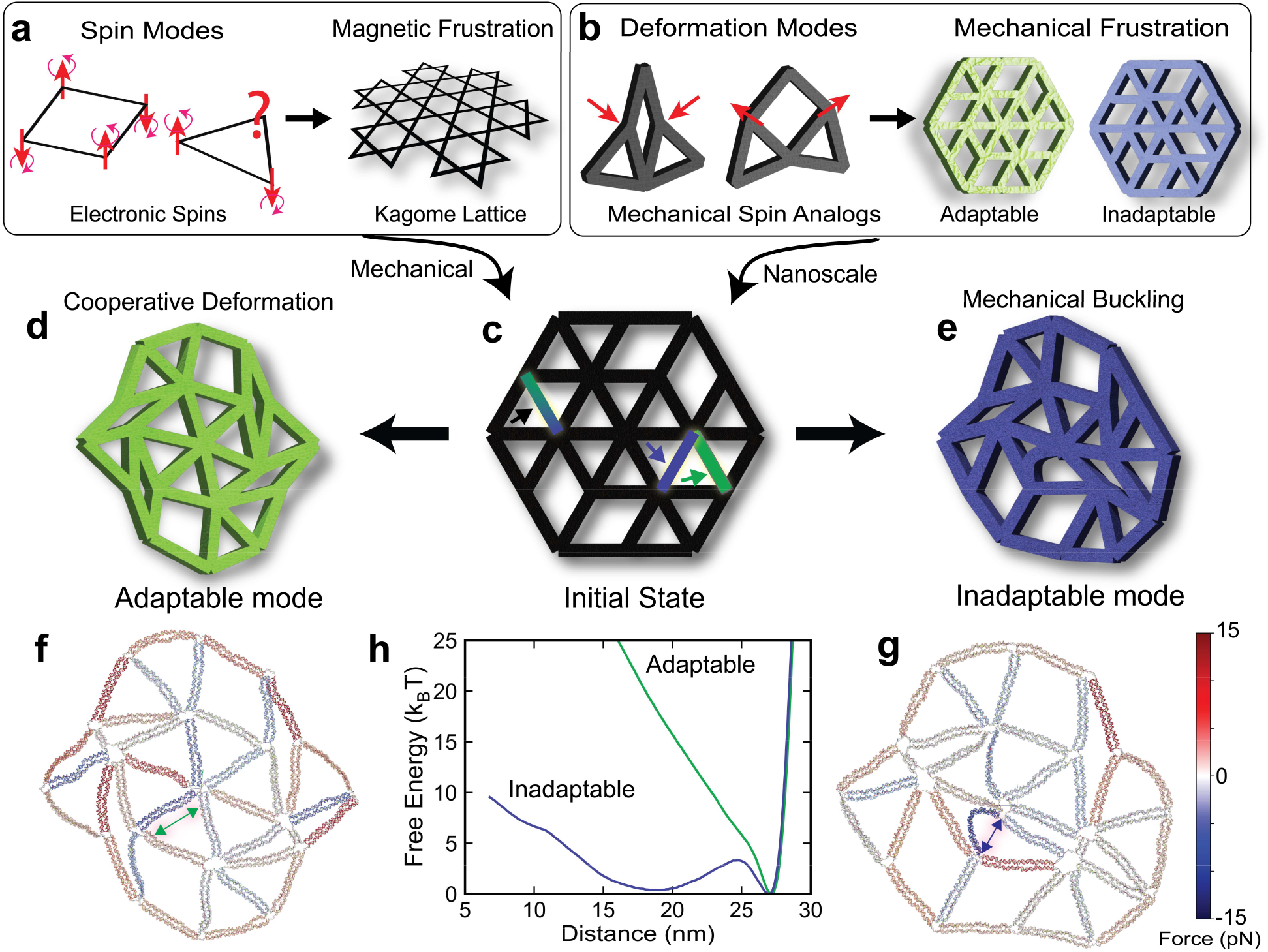
Geometric frustration in DNA origami. **a**, Interactions of electronic spins (red arrows) lead to a stable (ferromagnetic) state in a square arrangement, while a triangular design cannot minimize the energy (anti-ferromagnetic). The spins in the Kagome lattice exhibit complex magnetic behaviors. **b**, Electronic spin can be replaced by mechanical strain in two distinct deformation modes. Interactions between neighboring building blocks lead to adaptability (cooperative deformation) or inadaptability (frustration due to conflicting deformation). Both modes differ geometrically only in the orientation of a unit cell. **c**, Schematic of theoretical DNA origami design combining both modes into a single lattice, with edges made of 84-nt-long 2HB DNA. The edges highlighted in blue-green, blue, and green can serve as actuators (DNA jacks). **d** and **e**, Activating the blue-green edge in conjunction with green steers the structure into the adaptable state (**d**), while blue and blue-green together lead to the inadaptable state (**e**). **f** and **g**, MD simulated conformations of adaptable and inadaptable DNA. The structures are colored based on internal forces, with blue indicating compression and red indicating extension. **h**, Free energy profiles of adaptable and inadaptable modes using umbrella sampling. The x-axis is the distance of the edges indicated by green and blue arrows in **f** and **g**.

Buckling occurred consistently in a particular edge across multiple rounds of MD simulations. To delve deeper into the underlying mechanism dictating the observed conformations, we calculated the free energy profiles using umbrella sampling (see SI section S5 for computational details).^35^ Figure 1h presents the free energy landscapes of the two modes as a function of distance shown as green and blue arrows in Fig. 1f-g. The adaptable mode exhibits a steep-welled, single minimum in its free energy, while the inadaptable state shows two distinct minima: a gradual dip with a flat bottom from 17 to 20 nm and a minimum at ∼28 nm. The single minimum in the adaptable mode indicates that the most stable state occurs when the edge is fully extended, suggesting that strain is not concentrated at a particular point. Since all the edges in the state move coherently, the deformation of the cells and nodes is not impeded by any conflicts. Consequently, the corresponding free energy profile is similar to that of a single 2HB DNA edge (Fig. S11), implying that the deformation of the rest of the structure does not play a major role in affecting the free energy of the edge. In contrast, the dual minima of the inadaptable free energy landscape indicate two stable states where the edge is either buckled or fully extended. The short spike in free energy between the two minima shows that while the buckled and extended states are stable, intermediates that are partially bent are not. This hints that the edge experiences a snap buckling, i.e., the edge exists in either extended or buckled states, switching between them with sufficient energy overcoming the small activation barrier. The minimum at ∼28 nm implies that the buckling in the inadaptable state is transferred into other edges within the structure.

Unlike in macroscopic structures, random thermal fluctuations of a DNA metastructure increase the propensity of multistability, where the likelihood of other edges being stuck in buckled states also increases. Therefore, nanoscale frustrated metastructures exhibit a larger number of stable states in comparison to their macroscale counterparts. In addition, these states are not fixed as the inadaptable structure may transition from one stable mode to another by interchanging buckling. Figure 1f-g also indicates that while the adaptable state is symmetric, the inadaptable mode has a broken symmetry, suggesting that the distinct responses observed are caused by differences in symmetry as well. The inadaptable free-energy peak separating the two minima is relatively small (<5 *k*_*B*_*T* in magnitude), enabling transitions between the states and thus unlocking mechanical bistability at room temperature. The shape of the energy wells in Fig. 1h is a direct consequence of the design choice of using 2HB edges which are comparatively flexible. Choosing 4HB or 6HB edges will change the shape of the profile due to the higher energy cost associated with bending the stiffer edges, i.e., the wells of energy minima will become relatively deeper and narrower.^36^ The profiles can be tailored further, by changing the lengths of the ss-DNA sections at the vertices connecting the edges; in the current design, 60° and 120° junctions have 5 and 3 nt, respectively.

### Experimental demonstration

We demonstrated the lattice design experimentally using deformable DNA origami wireframes. First, we constructed DNA structures corresponding to both modes separately (see SI section S3 for experimental details). Starting from the synthesized initial undefined state (where the three adjustable edges are missing), we added two sets of staple strands to assemble either the adaptable or inadaptable modes (Fig. 2a). Figure 2b shows the theoretical designs, atomic force microscopy (AFM) scans, and contrast images of the adaptable state. As expected, the assembled DNA has mostly straight edges, with all interior triangles retaining shape. Each edge measures approximately 28 nm in length matching theoretical designs and roughly 4 nm in width, corresponding well with the expectation for 2HB DNA. In contrast, the inadaptable DNA displays evident buckling in some edges (highlighted by blue shades in the wireframe schematics in Fig. 2c), while the remaining edges are largely straight as designed. In rare cases, buckling occurs in multiple edges (first row). Both structures follow our designs accurately.

**Figure 2.**
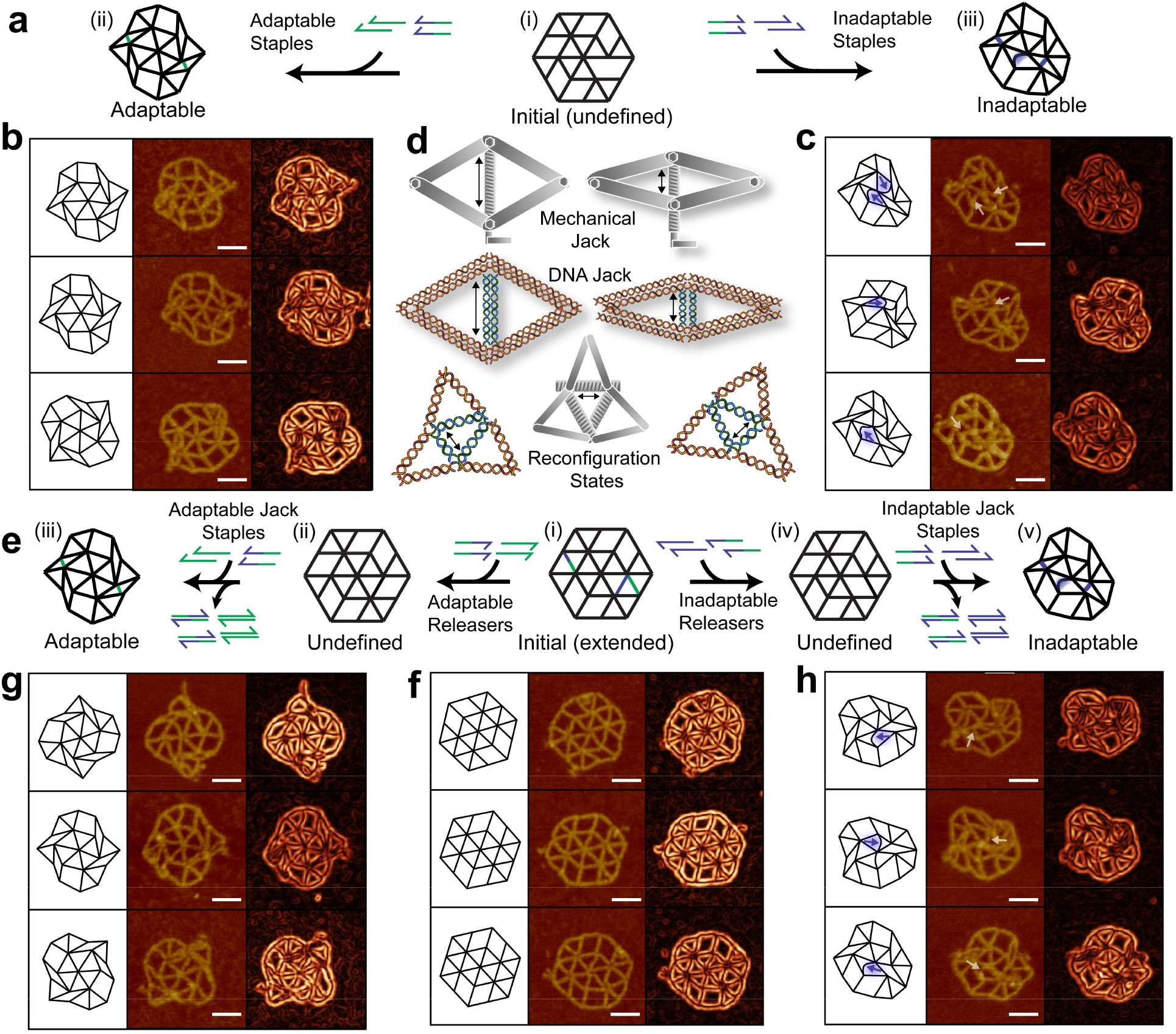
Experimental demonstration. **a**, Construction of adaptable and inadaptable structures from an initial undefined state by adding respective staples. **b** and **c**, Schematics, AFM scans, and contrast images of adaptable and inadaptable DNA nanostructures. The blue and white arrows highlight the buckled edge in the schematics and AFM images, respectively (**c**). **d**, Illustration of DNA jack edges analogous to a mechanical car jack and their function for external loading. By modulating the length of DNA jack edges, the nanostructure can be deformed into either adaptable or inadaptable states. **e**, Scheme for two-step DNA reactions for reconfiguration. This is achieved by first adding corresponding jack releaser strands to change from the initial extended mode to the undefined states and then introducing a set of jack staples to reconfigure into either inadaptable or adaptable modes. This chemical loading is reversible such that the origami can transition between initial, adaptable, and inadaptable states with appropriate inputs. **f**, Initial state with all three jack edges extended. **g** and **h**, Images of adaptable and inadaptable structures reconfigured after chemical loading. The scale bar in all AFM images is 50 nm.

Next, we studied deformation behaviors of our DNA metastructure upon external loading. Since it is difficult to apply forces precisely at specific vertices due to the small size of the structure, we used chemical loading to induce deformation. We achieved this by using adjustable DNA ‘jack’ mechanisms as illustrated in Fig. 2d. Like a car jack, the DNA jack can be adjusted in length via two-step DNA reactions – toehold-mediated strand displacement of one set of staples and re-annealing with a new set of staples of interest^30^. This actuation will deform the entire structure into different modes reversibly. Figure 2d (bottom) shows the DNA jack in action, as a part of the building block, increasing the number of possible deformation modes in comparison to the regular building block in Fig. 1c. In the deformation experiments, we began from the initial state with all jacks extended and deformed into either of the states by first adding the respective ‘releasers’ and then the staples for short jacks (Fig. 2e). The initial state with all jacks extended (Fig. 2f) transforms into either of the adaptable (Fig. 2g) or inadaptable (Fig. 2h) modes when subject to different loadings. The structures assumed in both states are consistent with those observed in the direct assemblies in Fig. 2b-c. The AFM scans of the intermediate steps during chemical loading are shown in Figs. S18 and S20. The lengths of the jack edges before and after reconfiguration change from 28 nm to <6 nm. The end-to-end distance of the buckled edges in the inadaptable state is about 18 nm, as predicted by free energy calculations in Fig. 1h. A considerable advantage of this chemical loading method is that it allows for reversible structural transformations between initial, adaptable, and inadaptable states (Figs. S22 and S23).

### Computational mechanics

We used coarse-grained MD models to study structural and mechanical behaviors during deformation. Compared to traditional elasticity theory predictions, MD simulations are advantageous as they account for both thermodynamic and mechanical properties. Instead of using chemical loading, simulations allow us to apply external forces directly to the DNA structure, enabling us thus to study how deformation and strain evolve over time. First, we removed the jack edges, leaving 10 bp short residues on each side to aid force application (highlighted by arrows in Fig. 3 a(i) and b(i)), and pulled them closer from 28 nm to 6 nm using harmonic traps in oxDNA (see SI for computational details). Figure 3 shows a series of snapshots representing effective (normalized) strain distributions in the adaptable and inadaptable modes. The strain map of the adaptable mode has a nearly uniform distribution throughout the domain. There are small segments of high strain (in red), which is a result of thermal fluctuations. No edges remain stuck in a state of excessive strain. Contrastingly, several edges of the inadaptable mode show significantly greater strain (and flexure) compared to their adaptable counterparts at the same timestep. While the strain fluctuates in most edges, it starts to develop in a particular edge and becomes more prominent with time (Fig. 3b (vii)-(xi)). Finally, strain is concentrated into the edge and the structure becomes stuck in a buckled state. Upon removal of the loading, both structures spontaneously return to their initial configurations (shown in (xii)). The average forces required to pull the structures is approximately 15 pN, while the maximum force that could be theoretically exerted is ∼57 pN (estimated from the binding free energies of jack staple sequences; see SI section S3 for details), suggesting that the chemical loading provided experimentally was sufficient^26^.

**Figure 3.**
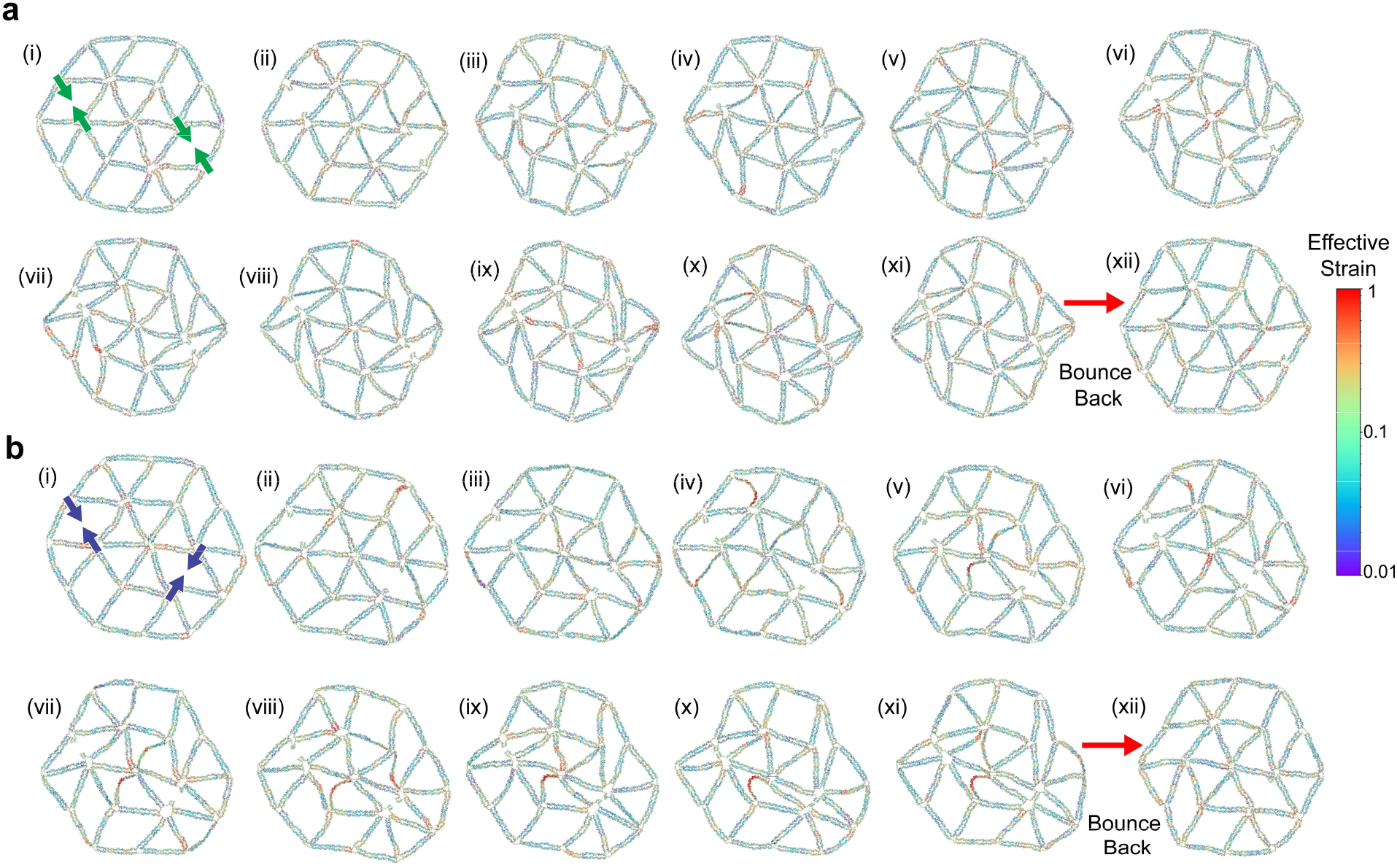
MD simulation of mechanical loading. **a**, Snapshots of oxDNA configurations under mechanical loading from (i) initial to (xi) adaptable states. Each snapshot represents the conformation in the intervals of 5×10^6^ steps. The edges being pulled are highlighted using green arrows in (i). The structure returns to the initial state after removal of external forces (xii). **b**, Snapshots of DNA conformations during deformation from (i) initial to (xi) inadaptable states. The inadaptable mode bounces back to the initial state (xii), with the buckled edge completely recovered. All the edges in the structures are colored based on normalized strain calculated using the local tangent vector 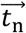 at each location. We use the tangent vector 5-nt upstream *(n-5)* and downstream *(n+5)* to obtain the strain at nt no. *n* as 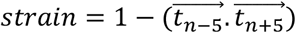 which ranges from 0 to 1. The buckled edge shows the highest local strain in **b** and small kinks also have relatively high strain in **a**.

Notably, most edges that show high strain in Fig. 3b are the diagonal struts of a rhombic group of edges similar in geometry to those shown in Fig. 2c and 2h. This is in fact an indicator of edges that are more likely to buckle. To understand this, we analyzed two (red and blue) rhombi each in the adaptable and inadaptable modes shown in the insets to Fig. 4a-b and plotted the bend deflection of the diagonal struts against the average angle enclosed during the simulation in Fig.3. Since there is no buckling in the adaptable mode, both the angle and bend deflection are nearly invariant throughout the deformation. Though the adaptable distributions deviate slightly (Fig. 4a (iii)), they return to an equilibrium angle of ∼60 degrees with minimal deflection. On the other hand, the distributions of the red and blue rhombi diverge in the inadaptable mode because the red diagonal edge is trapped into the buckled state during deformation while the blue edge remains extended (Fig. 4b (vi)-(x)).

**Figure 4.**
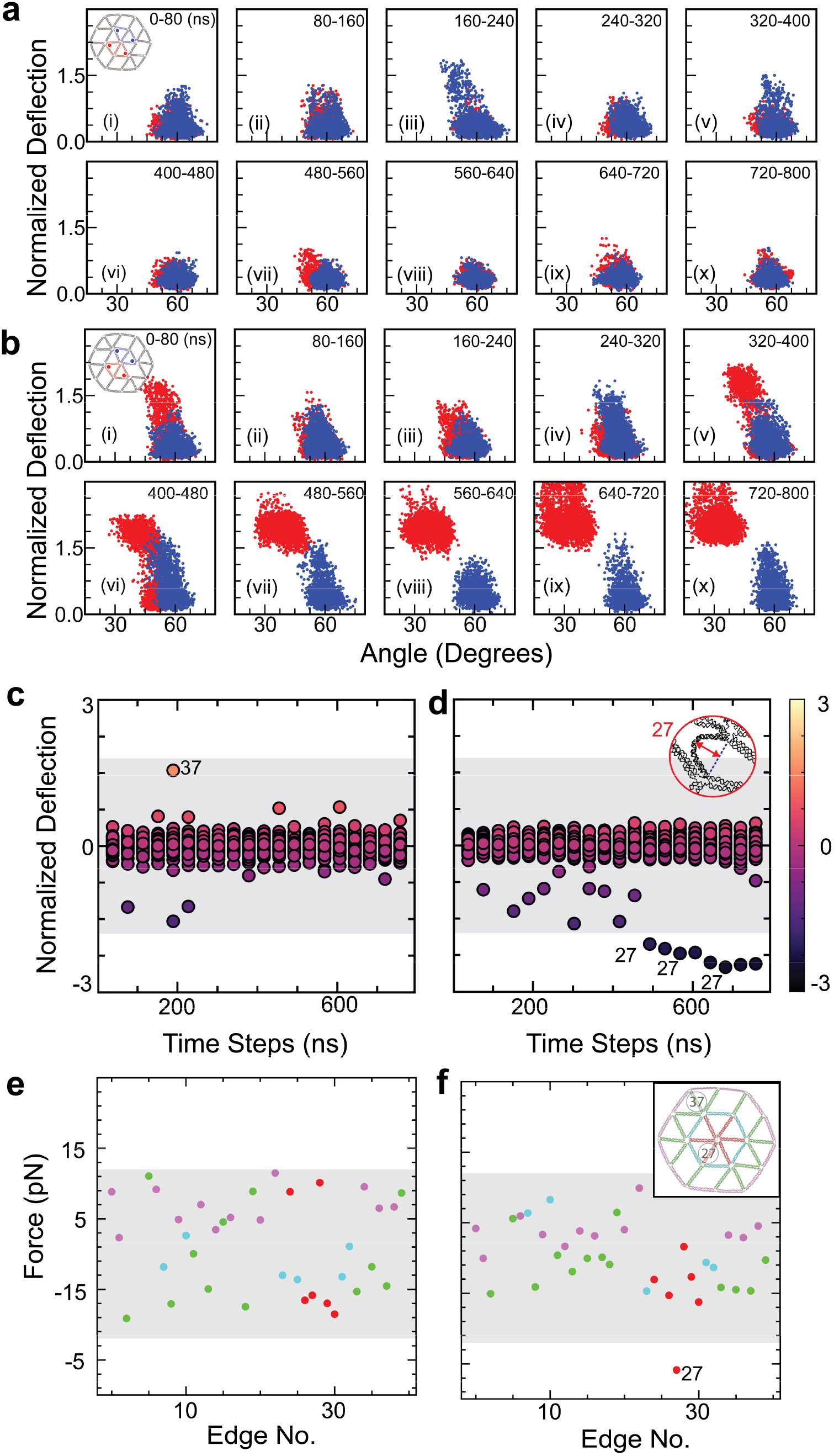
Structural and mechanical analysis. **a** and **b**, Bend deflection of central struts of colored rhombi (shown in red and blue in insets) vs average angle enclosed (red and blue dots in insets) for the adaptable (**a**) and inadaptable (**b**) states. Each colored dot represents the normalized deflection at a particular time instant in the given time interval. The snapshots in Fig.3 (i)-(x) are representative conformations for the intervals (i)-(x). While the adaptable state remains constant all throughout, the distributions of the inadaptable state diverge with time due to buckling of the red strut. Here, the bend deflection is the maximum distance from the line joining the ends (shown in **d** inset). **c** and **d**, Evolution of bend deflection of all edges with time for the adaptable (**c**) and inadaptable (**d**) modes. At each time instant, each circle represents the bend deflection of an edge. Two edges with relatively high bend deflection are numbered for reference. **e** and **f**, Internal forces for each edge extracted from MD simulations at equilibrium for adaptable (**e**) and inadaptable (**f**) states. The force distribution in **e** is more widespread, while most edges in **f** follow a narrow distribution with excessive compression concentrated in a single edge (edge 27). These patterns indicate the mechanism of strain distributions within the structure. The edges in **e** and **f** are also colored on the basis of relative position, highlighted by the inset in **f**. The edges experiencing significant flexure (37) and buckling (27) are highlighted. The complete numbering of all edges is given in Fig. S3.

Figure 4c-d shows the normalized bend deflection of each edge over time during deformation. With 36 edges in each structure, we plotted the average normalized bend deflection for each edge. Thus, each point represents the bend deflection of a particular edge at a particular instant of time. All edges of the adaptable mode shown in Fig. 4c stay within the confined region shaded gray, implying that none of the edges bend excessively. While few edges experience a flexure at some instants (marked edge 37), they bounce back and never permanently settle into a buckled state. However, the inadaptable mode has many more edges in the extrema of the gray region, with one of the edges experiencing severe buckling (marked edge 27 in Fig. 4d) during the latter half of the simulation. This supports the visual observations in Fig. 3b, with the buckled edge 27 having the highest strain. While this analysis focuses on planar effects of frustration, the structure in free solution likely experiences out-of-plane deformations with the edges buckling along the weakest direction (i.e., with the lowest bending modulus as shown in Figs. S12 and S13).

Examining the structures further, the final states of the simulation shown in Fig. 3 were used as input after ligating their jack edge residues. We simulated equilibrium conformations for a long period of time (i.e., 144×10^6^ steps). The average internal force at the central section of each edge was extracted for both modes (Fig. 4e-f). The edges were grouped based on their relative location within the structure to understand if that has any role in determining their stress states. They were grouped into outer hexagon (pink), inner hexagon (light blue), outer spokes (green), and inner spokes (red), as illustrated in the inset of Fig. 4f. When comparing the adaptable and inadaptable modes, the exterior hexagons are primarily under extension in both cases; however, the distribution of the inner hexagons differs markedly. In Fig. 4f, one edge (no. 27) is a clear outlier, experiencing greater compression relative to others and therefore buckling as seen in Fig. 3b. Figure 4f also indicates that the interior spokes are the edges most likely to buckle in the inadaptable state, with most being in compression. Additionally, the distribution of internal forces in Fig. 4e is wider, suggesting an equal spread of forces throughout the structure. Conversely, most edges of the inadaptable state in Fig. 4f lie in a narrow distribution with the buckled edge being a lone outlier, suggesting localization of stresses. External loading and internal force simulations together elucidate stark differences in response of the two modes.

## DISCUSSION

Our work showcases the powerful synergy between DNA self-assembly techniques and mechanical design principles, paving the way to the development of a previously unexplored class of nanoscale frustrated metamaterials. By leveraging proper geometric design, we were able to tune the free-energy profiles of nanostructures and precisely tailor their response to external stimuli. DNA origami makes the metastructures more versatile by engineering both adaptability and inadaptability into a single lattice, which is difficult to achieve with pure top-down approaches.

The ability to route strain from external loading can be leveraged to develop methods of storing mechanical energy inside nanostructures. In nature, motor proteins like kinesin use controlled storage and release of strain energy to achieve directional motion with high processivity and directionality^37, 38^. The extension and contraction of bundles of actomyosin filaments inside a cell are crucial for enabling cell motility^39^. The design approaches described here may be extended to develop synthetic motors capable of localized storage and release of strain on-demand. This feature could also be used to perform nanoscale mechanical computations through shape memory. For example, different sequences of input stimuli would lead to unique strain states, potentially enabling a retrieval of the sequence of inputs based on the final strain state alone.

Frustration plays a crucial role in several biomolecular functions like protein folding, allostery, catalysis, and DNA-protein binding^40^. Allostery is often triggered by the transfer of strain from one part of a protein to another upon ligand binding at a given site, or other external stimuli. Although chemically different, the mechanics underlying this phenomenon are similar to those engineered in this work, namely, the strategic redistribution of strain through chemo-mechanical coupling. Replicating allosteric phenomena in synthetic systems has been challenging so far, but DNA nanotechnology offers an ideal platform for their construction and study. Although the present study focused on 2D deformations, it can serve as a steppingstone toward extending these principles to the design of 3D metastructures.

While the current design exhibited dual energy minima, further modifications could help engineer more complex energy landscapes with multiple minima. The mechanical response of the metastructure can be further controlled by changing mechanical parameters such as edge thickness and number of nicks within the same design geometries. The concept of mechanical frustration can thus be employed to provide a diverse set of design tools for programming both free energy and mechanical properties. As shown here, the tools provided by structural DNA nanotechnology are well suited to implement mechanical frustration at the nanoscale and could potentially be used to understand natural systems as well as enhance the capabilities of architected nanomaterials.

## ACKNOWLEDGMENTS

The authors thank Prof. P. Sulc, Dr. M. Matthies, and M. Sample at Arizona State University for help with free energy calculations and Prof. A. Arrieta at Purdue University for insightful discussion. This work was financially supported by the U.S. Department of Energy, Office of Science, Basic Energy Sciences under award no. DE-SC0020673. J.H.C. also acknowledges the Friedrich Wilhelm Bessel Research award from the Alexander von Humboldt Foundation.

## REFERENCES

1. Zhang, H., Guo, X., Wu, J., Fang, D. & Zhang, Y. Soft Mechanical Metamaterials with Unusual Swelling Behavior and Tunable Stress-Strain Curves. Science Advances 4, eaar8535 (2018).

2. Hwang, D., Barron, E.J., Haque, A.B.M.T. & Bartlett, M.D. Shape Morphing Mechanical Metamaterials through Reversible Plasticity. Science Robotics 7, eabg2171 (2022).

3. Lakes, R. Foam Structures with a Negative Poisson’s Ratio. Science 235, 1038–1040 (1987).

4. Mintchev, S., Shintake, J. & Floreano, D. Bioinspired Dual-Stiffness Origami. Science Robotics 3, eaau0275 (2018).

5. Faber, J.A., Arrieta, A.F. & Studart, A.R. Bioinspired Spring Origami. Science 359, 1386–1391 (2018).

6. Kapnisi, M. et al. Auxetic Cardiac Patches with Tunable Mechanical and Conductive Properties toward Treating Myocardial Infarction. Advanced Functional Materials 28, 1800618 (2018).

7. Smith, D.R., Pendry, J.B. & Wiltshire, M.C. Metamaterials and Negative Refractive Index. Science 305, 788–792 (2004).

8. Cummer, S.A., Christensen, J. & Alù, A. Controlling Sound with Acoustic Metamaterials. Nature Reviews Materials 1, 1–13 (2016).

9. Kasuya, T. A Theory of Metallic Ferro-and Antiferromagnetism on Zener’s Model. Progress of theoretical physics 16, 45–57 (1956).

10. Kanô, K. & Naya, S. Antiferromagnetism. The Kagomé Ising Net. Progress of theoretical physics 10, 158–172 (1953).

11. Meeussen, A.S., Oğuz, E.C., Shokef, Y. & Hecke, M.v. Topological Defects Produce Exotic Mechanics in Complex Metamaterials. Nature Physics 16, 307–311 (2020).

12. Udani, J.P. & Arrieta, A.F. Taming Geometric Frustration by Leveraging Structural Elasticity. Materials & Design 221, 110809 (2022).

13. Meeussen, A.S., Oğuz, E.C., Hecke, M.v. & Shokef, Y. Response Evolution of Mechanical Metamaterials under Architectural Transformations. New Journal of Physics 22, 023030 (2020).

14. Rothemund, P.W.K. Folding DNA to Create Nanoscale Shapes and Patterns. Nature 440, 297–302 (2006).

15. Zhang, F. et al. Complex Wireframe DNA Origami Nanostructures with Multi-Arm Junction Vertices. Nature Nanotechnology 10, 779–784 (2015).

16. Wintersinger, C.M. et al. Multi-Micron Crisscross Structures Grown from DNA-Origami Slats. Nature Nanotechnology 18, 281–289 (2023).

17. Liu, L., Li, Z., Li, Y. & Mao, C. Rational Design and Self-Assembly of Two-Dimensional, Dodecagonal DNA Quasicrystals. Journal of the American Chemical Society 141, 4248–4251 (2019).

18. Iinuma, R. et al. Polyhedra Self-Assembled from DNA Tripods and Characterized with 3d DNA-Paint. Science 344, 65–69 (2014).

19. Song, J. et al. Reconfiguration of DNA Molecular Arrays Driven by Information Relay. Science 357, eaan3377 (2017).

20. Marras, A.E., Zhou, L.Su, H.-J. & Castro, C.E. Programmable Motion of DNA Origami Mechanisms. Proceedings of the National Academy of Sciences 112, 713–718 (2015).

21. Zhou, L., Marras, A.E.Su, H.-J. & Castro, C.E. DNA Origami Compliant Nanostructures with Tunable Mechanical Properties. ACS Nano 8, 27–34 (2014).

22. Shi, X. et al. A DNA Turbine Powered by a Transmembrane Potential across a Nanopore. Nature Nanotechnology 19, 338–344 (2024).

23. Kopperger, E. et al. A Self-Assembled Nanoscale Robotic Arm Controlled by Electric Fields. Science 359, 296–301 (2018).

24. Büchl, A. et al. Energy Landscapes of Rotary DNA Origami Devices Determined by Fluorescence Particle Tracking. Biophysical Journal 121, 4849–4859 (2022).

25. Kim, M. et al. Harnessing a Paper-Folding Mechanism for Reconfigurable DNA Origami. Nature 619, 78–86 (2023).

26. Li, R., Chen, H. & Choi, J.H. Auxetic Two-Dimensional Nanostructures from DNA. Angewandte Chemie International Edition 60, 7165–7173 (2021).

27. Li, R., Madhvacharyula, A.S., Du, Y., Adepu, H.K. & Choi, J.H. Mechanics of Dynamic and Deformable DNA Nanostructures. Chemical Science 14, 8018–8046 (2023).

28. DeLuca, M. et al. Thermally Reversible Pattern Formation in Arrays of Molecular Rotors. Nanoscale 15, 8356–8365 (2023).

29. Wang, Y. et al. Steric Communication between Dynamic Components on DNA Nanodevices. ACS Nano 17, 8271–8280 (2023).

30. Li, R., Chen, H. & Choi, J.H. Topological Assembly of a Deployable Hoberman Flight Ring from DNA. Small 17, 2007069 (2021).

31. Chhabra, H. et al. Computing the Elastic Mechanical Properties of Rodlike DNA Nanostructures. Journal of Chemical Theory and Computation 16, 7748–7763 (2020).

32. Yoo, J. & Aksimentiev, A. In Situ Structure and Dynamics of DNA Origami Determined through Molecular Dynamics Simulations. Proceedings of the National Academy of Sciences 110, 20099–20104 (2013).

33. Poppleton, E., Romero, R., Mallya, A., Rovigatti, L. & Šulc, P. Oxdna.Org: A Public Webserver for Coarse-Grained Simulations of DNA and Rna Nanostructures. Nucleic Acids Research 49, W491–W498 (2021).

34. Rovigatti, L., Šulc, P., Reguly, I.Z. & Romano, F. A Comparison between Parallelization Approaches in Molecular Dynamics Simulations on Gpus. Journal of Computational Chemistry 36, 1–8 (2015).

35. Torrie, G.M. & Valleau, J.P. Nonphysical Sampling Distributions in Monte Carlo Free-Energy Estimation: Umbrella Sampling. Journal of Computational Physics 23, 187–199 (1977).

36. Wong, C.K., Tang, C., Schreck, J.S. & Doye, J.P.K. Characterizing the Free-Energy Landscapes of DNA Origamis. Nanoscale 14, 2638–2648 (2022).

37. Carter, N.J. & Cross, R.A. Mechanics of the Kinesin Step. Nature 435, 308–312 (2005).

38. Rosenfeld, S.S., Fordyce, P.M., Jefferson, G.M., King, P.H. & Block, S.M. Stepping and Stretching: How Kinesin Uses Internal Strain to Walk Processively. Journal of Biological Chemistry 278, 18550–18556 (2003).

39. Bao, G. Mechanics of Biomolecules. Journal of the Mechanics and Physics of Solids 50, 2237–2274 (2002).

40. Ferreiro, D.U., Komives, E.A. & Wolynes, P.G. Frustration in Biomolecules. Quarterly Reviews of Biophysics 47, 285–363 (2014).

