## Supplementary Information for "Realizing Mechanical Frustration at the Nanoscale Using DNA Origami"

**Table of Contents**

S1 Design of Geometric Frustration
S2 Materials and Methods
S3 DNA Origami Design
S4 OxDNA Simulations
S5 Free Energy Calculations
S6 Bending and Out-of-plane Deformation
S7 Additional AFM Images
S8 DNA Sequences
S9 References

### S1 Design of Geometric Frustration

Geometric frustration arises from the inability of a system to minimize its energy due to competing interactions. It commonly arises from interactions between electronic spins and has been studied within the framework of the Ising spin model and the Kagome lattice. Utilizing the geometry of the 2D Kagome lattice, similarly shaped mechanical building blocks can be developed in which electronic spin is replaced with mechanical deformation[1] that gives rise to geometric frustration. The building block in this design has two deformation modes analogous to upward and downward electronic spins as seen in Figs. 1b and S1a. Arranging these blocks into a hexagonal arrangement allows interactions between neighboring blocks. Fig. S1b illustrates the deformations of the building blocks for adaptable mode with all directions of deformations in synchrony. The inadaptable mode due to conflicting deformations (at the node highlighted in Fig. S1c) exhibits geometric frustration due to opposing deformations of neighboring building blocks. The deformations are dictated by the points where the structure is actuated by external loading. As will be discussed in the main text and in further detail in Section 3, the actuation of the structure using reconfigurable struts (jack edges) enables the fusion of both adaptability and inadaptability into a single metastructure. The differences between both structures are minimal – the only difference being orientation of the building block on the bottom right. This framework may be expanded to other 2D planar lattices, with a variety of deformation modes, by using a combinatorial approach for arrangement of building blocks.

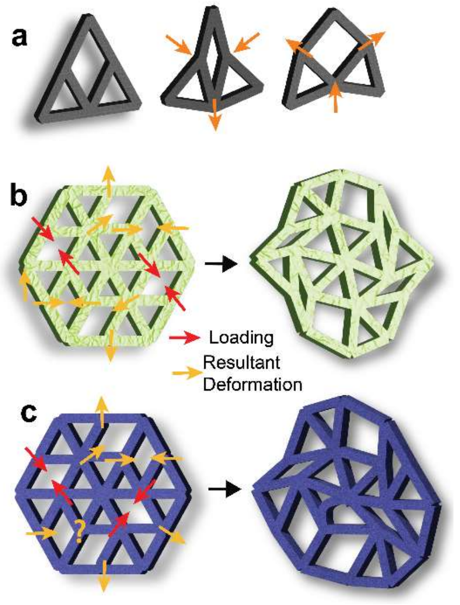


**Figure S1.** Representative diagrams for assembling building blocks into adaptable and inadaptable geometries. (**a**) A triangular building block and its two basic deformation modes. (**b**) Loading and resultant deformations for the adaptable mode. External forces are applied at the arrows colored in red. (**c**) Loading and deformations mapped for the inadaptable mode. The external forces applied deform the structure in conflicting directions, leading to frustration. The node highlighted with a question mark is the node of conflict.

### S2 Materials and Methods

#### S2.1 DNA origami assembly

All DNA oligomers were purchased from Integrated DNA Technologies (IDT). The M13mp18 scaffold used was procured from Bayou Biolabs. All the other chemicals used were purchased from Sigma Aldrich. DNA origami nanostructures were constructed by mixing 5 nM M13 scaffold strands with 4× DNA staple oligonucleotides for regular edges and 8× DNA staples for jack (reconfigurable) edges in 1× TAE buffer (solution of 40 mM trisaminomethane 1 mM ethylenediaminetetraacetic acid (EDTA) disodium salt, and 20 mM acetic acid) with 6 mM magnesium acetate (referred to as TAEM6). This mixture was annealed from 95 to 65 °C at 1 °C per 2 mins, 65 to 60 °C at 1 °C per 25 mins, 60 to 50 °C at 1 °C per 60 mins, and 50 to 35 °C at 1 °C per 40 mins after which it was cooled down to 4 °C. The duration of the complete thermal cycle was about 1 day. The adaptable and inadaptable structures shown in Fig. 2b-c were constructed by adding respective staples for two short jack edges and third extended jack (Fig. 2a (ii) and (iii)), along with staples for other edges. The initial extended state (Figs. 2e (i) and 2f) was prepared by adding the extended jack staples with toeholds for three jack edges along with staples for other edges. These mixtures were subjected to the same thermal cycle mentioned above.

#### S2.2 Chemical loading via two-step DNA reactions

The initial state prepared was the starting point for reconfiguration to both adaptable and inadaptable states. This was achieved by two-step reactions: toehold-mediated strand displacement with releasers and re-annealing with a new set of staples. After the synthesis of the initial state, the excess staples were removed using 100 kDa Pall Nanosep centrifugal filters, by centrifuging a mixture (55 µL) of DNA origami solution with 400 µL of TAEM6 buffer at 5000 rpm for 3 mins. This process was repeated twice more after which the origami was redispersed in new TAEM6 buffer. The releasers of the respective jack edges depending on the desired final state (adaptable/inadaptable) were added in 4× concentration to the purified mixture and incubated for ~14 hrs (55 °C for 3 hrs and then cooled down to 35 °C over 11 hrs). The resulting origami solution (containing adaptable or inadaptable undefined states) was again purified to remove excess staples. Then, selected short jack staples were added to the undefined states and incubated for 16 hrs (55°C for 4hrs and cooled slowly to 35 °C over 12 hrs) to deform the structure into the final configuration. The resulting DNA was again purified by centrifugation to remove excess staples and prepare the sample for AFM imaging.

Reversibility between adaptable and inadaptable states was tested by starting from either adaptable or inadaptable states, adding releaser staples, and incubating for ~14 hrs (55°C for 3 hrs and then cooled down to 35 °C over 11 hrs). The initial undefined state was then obtained (with all three jack edge staples missing). After purification to remove excess staples, respective short jacks were added to the initial undefined state to reach either adaptable or inadaptable state based on the reconfiguration direction. The AFM scans before and after reconfiguration showed buckled edges recovering completely indicating that the structures did not experience permanent deformations.

#### S2.3 AFM Imaging

AFM imaging was performed in air using the Peak-Force Tapping mode using a Bruker Dimension Icon with SCANASYST-AIR probes. The DNA origami samples were diluted to 0.5-1 nM during deposition. The samples were prepped for AFM by adding 2 µL of DNA solution and 8 µL of TAEM6 buffer, which was incubated for 5 mins. To improve adhesion of DNA to mica, 20 µL buffer with 2.5 mM $NiCl_2$ was added and allowed to incubate for an additional 2 mins. The sample was then blown dry with compressed air and washed with 80 µL deionized water. Each mica sample was scanned at multiple locations using typical scan sizes of 5 × 5 µm, 2 × 2 µm, and 500 × 500 nm.

**S3 DNA Origami Design**

The wireframe DNA origami was designed using Cadnano[2] and Scadnano. [3] The structure consisted of 38 2HB edges on a hexagonal lattice, each edge being 84 nucleotides long (~28 nm in length and ~4 nm in diameter). Three of these edges (i.e., 3 jacks) were designed to be reconfigurable with staples having additional toeholds (8 nts) and a slightly different arrangement for maximum contraction in length. By design, all staples were 42-nt long except the jack staples which had lengths of 50 or 20 nts (including the 8-nt toehold). The unused free scaffold segment was concentrated into loops on two edges which also served as indicators of the orientation of the structure during AFM. The Scadnano design of the structure is shown in the Fig. S2. The single-stranded nucleotide (ssnt) joints within the structure were designed following recommendations by a previous work on wireframe DNA origami by Yan and coworkers.[4] Joints with an expected angle of 120° have 3 ssnt, while 5 ssnts were used for a desired angle of 60° (Fig. S3 inset). The nomenclature for the staple sequences is based on their position following the convention of first going left to right and then top to bottom. The edges are also assigned numbers from 0 to 38, which are later used during analysis. The DNA structure as well as their numbers assigned to each edge are shown in Fig. S3. The edges no. 4, 25, and 17 are the reconfigurable jack edges. The jack staple arrangements and the two-step DNA reconfiguration for chemical loading is shown in Fig. S4. NUPACK[5] analysis estimates the free energy $\Delta G$ to be ~171.3 kcal/mol per jack (gained from the addition of short jacks). The $\Delta x$ (i.e., the change in length after contraction) is calculated to be about 20 nm. The maximum force that can be applied theoretically during chemical loading is therefore ~57 pN ($\Delta G/\Delta x$). This is well beyond the actual forces required (obtained from MD simulations) that are in the range of approximately 15 pN (Fig. S5). Thus, the design ensures that adequate energy is provided to the system for chemical loading. The extended jack edges (Fig. S4a) were shortened by first adding releasers to remove all the jack staples (Fig. S4b) and then adding new jack staples which bound to each contiguous scaffold segment, pulling the ends together from ~28 nm in the initial state to approximately 5 nm in the final state (Fig. S4c)

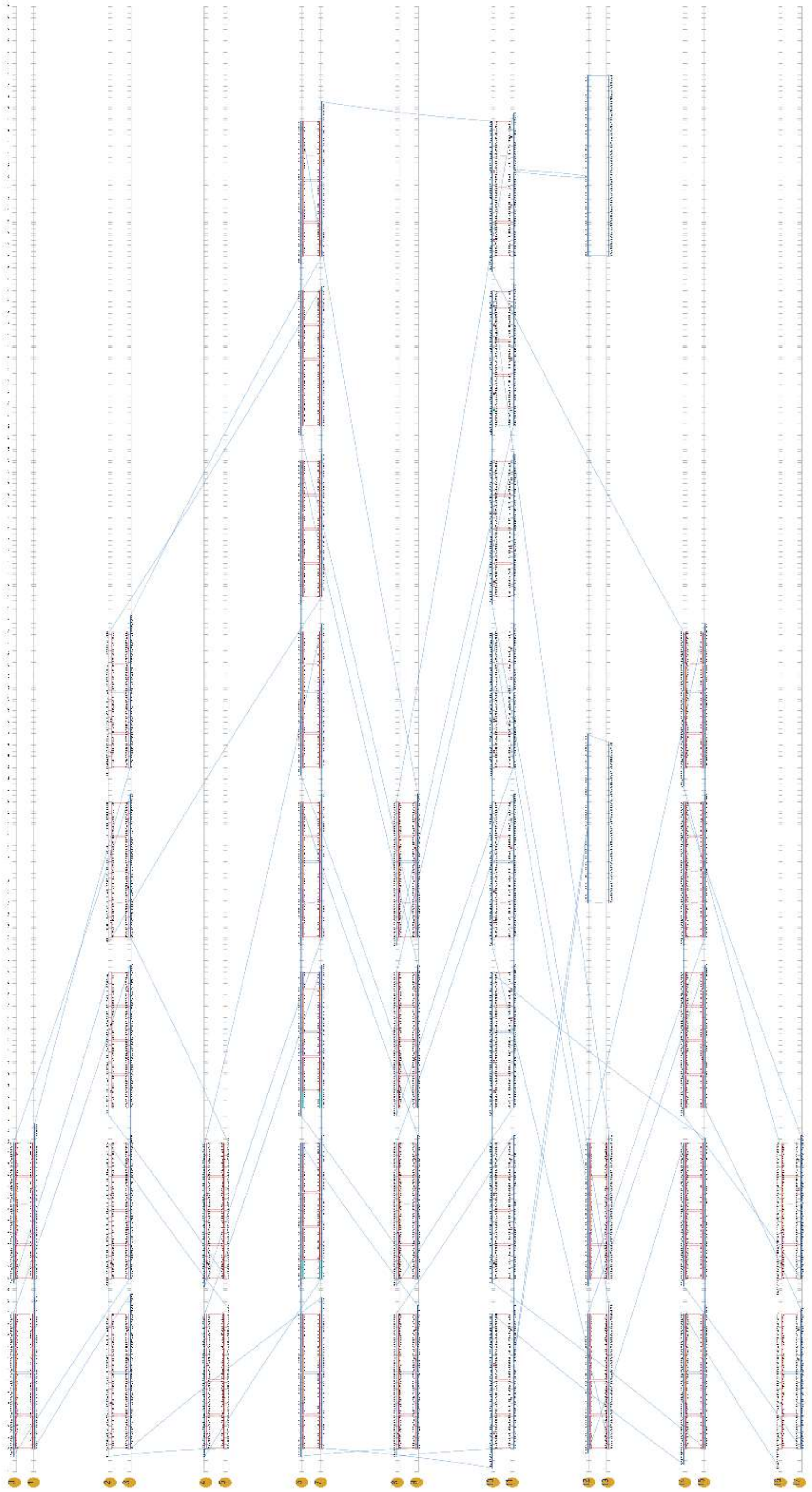

**Figure S2.** Layout of the theoretical origami design in Scadnano.

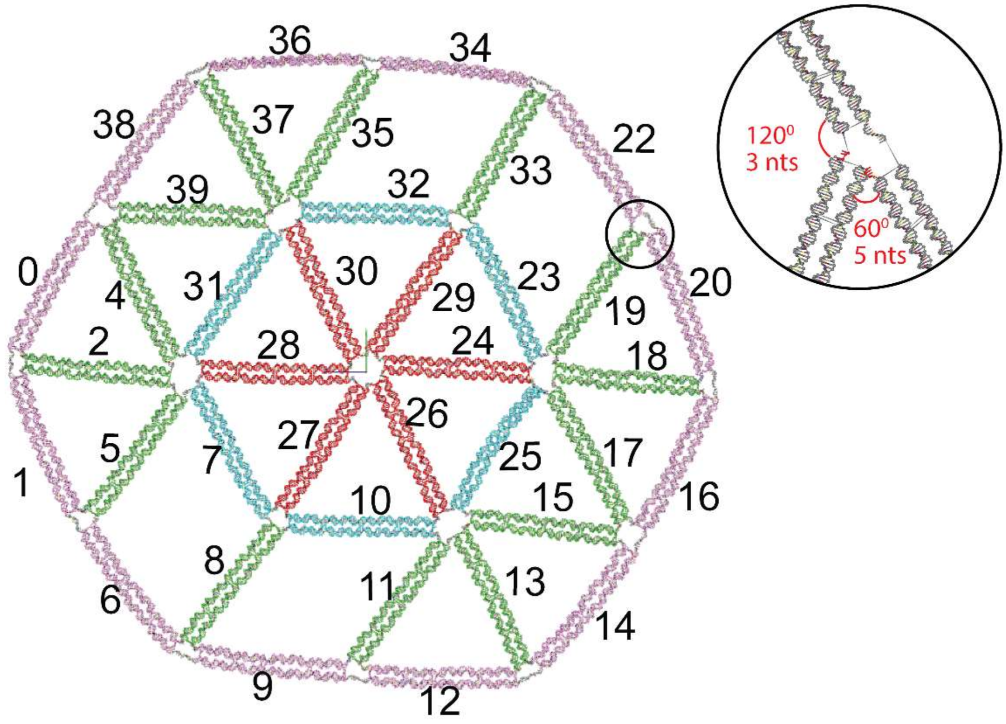


**Figure S3.** Metastructure design labelled with assigned edge numbers. The edges 4, 25, and 17 are jack edges that are used selectively to apply chemical loading. These edges are also color coded based on their relative position as described in Fig. 4 in the main text. The number of free ssnts are determined by the vertex angle and were fixed to be 3 nts for 120° and 5 nts for 60° (inset).

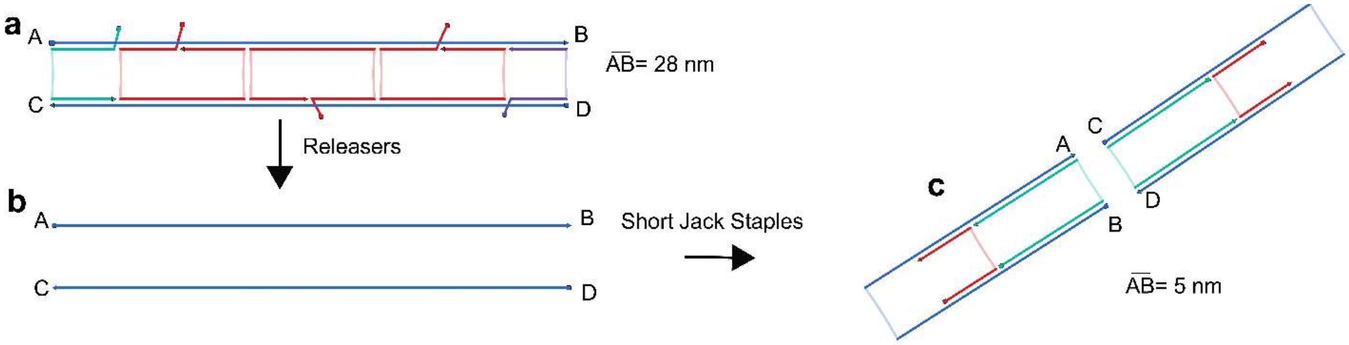


**Figure S4.** Staple arrangements in jack edges during strand displacement. (**a**) Extended jack edge having staples equipped with additional toeholds. (**b**) Free scaffold segments exposed after removal of staples due to addition of releasers. (**c**) Contraction in length after addition of short jack staples. The jacks are contracted by squeezing scaffold segments AB and CD from an initial length of 28 nm to 2HB loops of 5 nm in width.

**S4 OxDNA Simulations**

We studied our DNA metastructure with coarse-grained MD simulations using the oxDNA[6-8] platform (on a NVIDIA GTX 2060 GPU). All simulations performed used average sequence parameters at a temperature of 300 K with a diffusion coefficient of 2.5 and a salt concentration of 0.5 M.[9] The time increment *dt* was 0.005 (approximately 15 fs). The equilibrium structure of the initial state was obtained by first relaxing the Cadnano structure with greater backbone force for $10^5$ steps after which MD simulations were performed for $6\times10^6$ steps. Parts of the reconfigurable edges of the equilibrium conformation in the initial state were deleted using oxView to prepare the input files for mechanical deformations. A few base paired nucleotides (10 or 11 bps) were left uncut to prevent the 2HB helices from fraying when applying external loading using harmonic traps available in oxDNA. The jack edge residues were subject to spring forces, moving at a constant velocity and covering about 20 nm over $50\times10^6$ steps. We grouped the nucleotides in the oxDNA configuration and topology files were first grouped based on their edges with DBSCAN clustering for ease of analysis. The edge numbers are consistent with those shown in Fig. S3.

**S4.1 Mechanical deformation**

The harmonic traps exerted nearly constant and equal forces on both jacks. The traps started from the initial positions of the jack ends and moved at a rate of ~$2.4\times10^{-7}$ nm per step towards each other. The force exerted by the trap can be calculated by

$$\overrightarrow{F_{trap}} = k(\overrightarrow{r_{trap}} - \overrightarrow{r_{jack-nts}})$$

where $k$ is the stiffness set to 0.1 in the simulation (i.e., ~5.7 pN/nm). $\overrightarrow{r_{trap}}$ and $\overrightarrow{r_{jack-nts}}$ are the instantaneous position vectors of the trap and jack edge nts, respectively. The evolution of external loading with time in both modes is shown in Fig. S5. The slight increase in forces in the end is when both jack ends come in contact and are pulled past each other. Both final states after loading return to their initial conformations upon removal of external forces, implying that these structures can potentially be used to store mechanical energy. The trajectory as well as the average jack distance for the adaptable and inadaptable modes on removing external forces is shown in Figs. S6 and S7, respectively. The separation between the jack residues is gradual for the adaptable mode. The inadaptable mode experiences relatively a rapid recovery, with fluctuations later in the simulations. Therefore, both structures release stored energy in different ways.

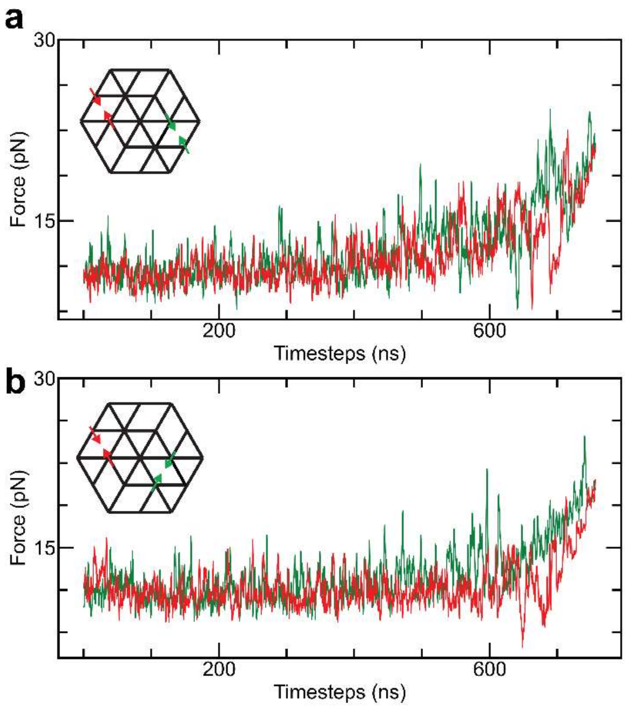


**Figure S5.** Forces exerted by harmonic traps for mechanical loading. The average force exerted by the harmonic traps at each jack edge of the adaptable (**a**) and inadaptable (**b**) modes. Each jack location is plotted with a different color as shown in the inset.

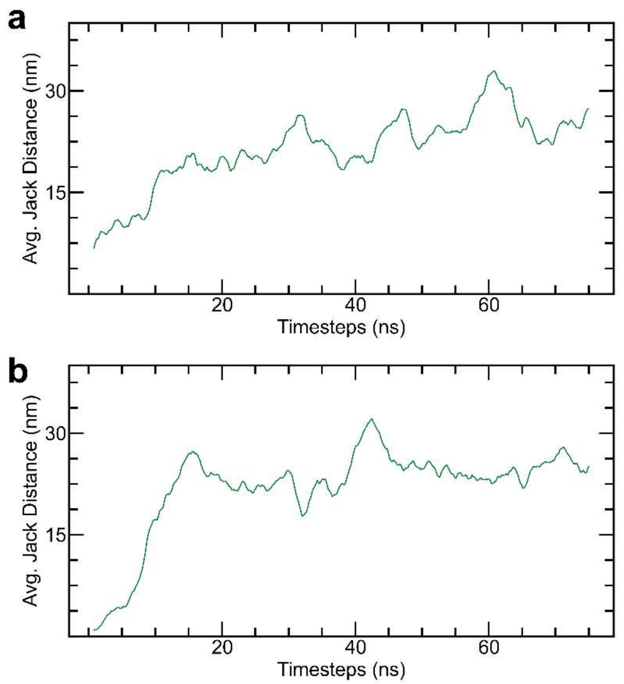


**Figure S6.** Evolution of end-to-end jack distance during recovery after removal of external forces. The average jack separations with time for the adaptable (**a**) and inadaptable modes (**b**). While the adaptable structure maintains a gradual increase all throughout, a sudden spike followed by a gradual increase is observed in the inadaptable mode. This highlights the differences in the way the energy is stored between adaptable (dispersed) and inadaptable (localized) modes.

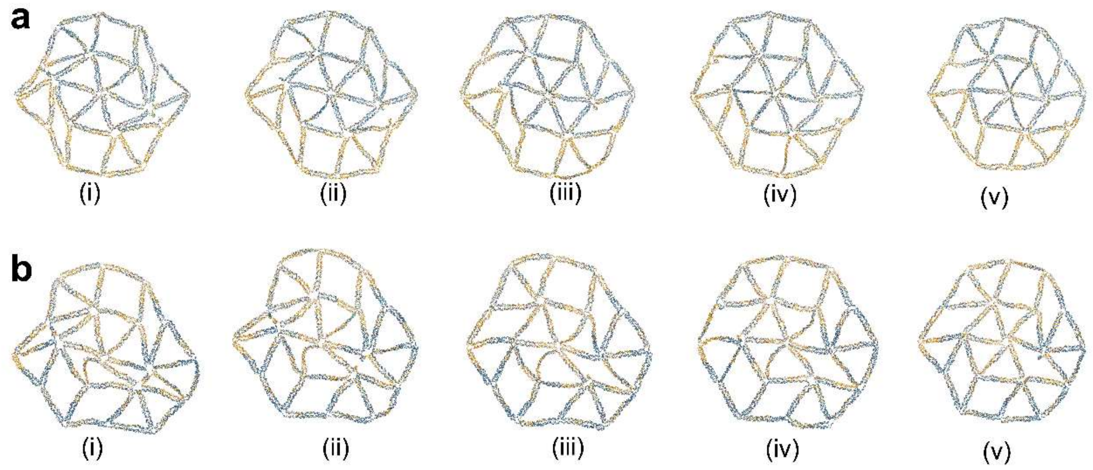


**Figure S7.** Snapshots of adaptable (**a**) and inadaptable structures (**b**) after removal of external loading in increments of $1.2 \times 10^6$ steps ((i) through (v)). Both structures return to their initial configurations after roughly $6 \times 10^6$ steps.

### S4.2 Bend deflection analysis

The bending of each edge during the simulation was quantified to understand mechanics and discern buckling from thermal fluctuations. For each edge in the structure, an averaged configuration was obtained by taking the average of center of masses of the 4 nt at each section of the 84-nt-long 2HB edge (i.e., 84 sections per edge) as illustrated in Fig. S8. The bend deflection was defined as the maximum distance of the averaged curve for each edge from the straight line (displacement) joining the edges. The calculations were then normalized by dividing with 4 nm (diameter of the 2HB edge) to prevent thermal fluctuations from being classified as bending. The normalized bend deflection values ranged from -3 to 3. The bend deflection vs time plot in Fig. 4c-d was obtained by averaging bend deflection values over windows of $500\times10^3$ steps for each of the 36 edges in each mode. Note the number of edges is not 38 but 36, since two reconfigurable jacks are partially cut off for each mode.

#### S4.3 Local strain visualization

A quantitative estimate for the bending/strain was developed by defining the local strain as a function of the dot product of the tangent vectors at different sections. The strain was obtained by using the formula

$$strain = 1 - (\overrightarrow{t_{n-5}} \cdot \overrightarrow{t_{n+5}}),$$

where $\overrightarrow{t_{n-5}}$ and $\overrightarrow{t_{n+5}}$ are tangent vectors, 5 nts upstream and downstream of the section where strain is being calculated. This was computed at multiple sections in each edge at a particular time instant. This method is illustrated in Fig. S8. Given the formula used, the first and last 5 sections of the edge are excluded from calculations. Therefore, each edge has 74 data points of strain. The plots are colored using a 'jet' colormap in oxView.

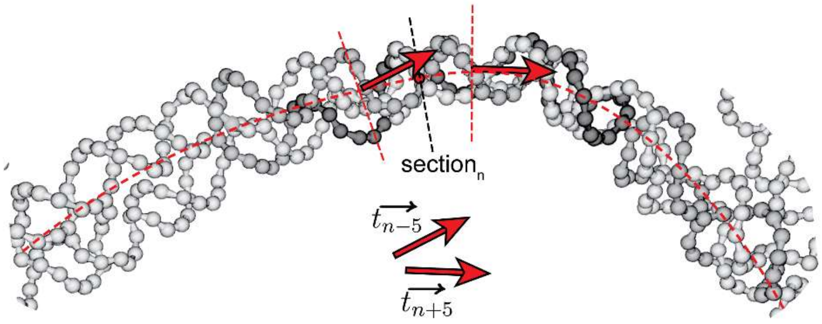


**Figure S8.** Calculation of local strain. Two unit tangent vectors upstream and downstream of the section of interest. The dot product of these vectors is subtracted from 1 to ensure higher strain has values closer to 1 while lower strain has values closer to 0, in line with the traditional definition of mechanical strain.

#### S4.4 Estimation of inter-nucleotide forces

Estimating internal forces can explain the differences in mechanical response of both modes. A prior work by Engel. et. al.[10] outlines a method to extract reliable estimates of inter-nucleotide forces in DNA origami. The tension (or compression) at a particular section of an origami edge can be obtained by summing up the forces acting on all the nucleotides upstream of the section plane exerted by all the nucleotides that are downstream of the section plane (Fig. S9). Although instantaneous magnitudes of this sum can be large, the average over a long timeframe results in realistic estimates of the inter-nucleotide forces. The input conformations for these simulations were generated by ligating the jack cutoff ends after being pulled close together in the loading simulations described earlier. The newly ligated structures were equilibrated for $6\times10^6$ steps. After equilibration, each structure was simulated for an additional $144\times10^6$ steps to estimate internal forces. The forces were computed at the central section of each edge (36 edges in total). The dot product of the computed net force vector and the local tangent vector at the central section yielded the net compression/tension. For compressive forces the dot product was negative while it remained positive for tension. The calculated force estimates range between -10 to 10 pN. The buckled edge in the inadaptable state stood out highlighting excessive compression.

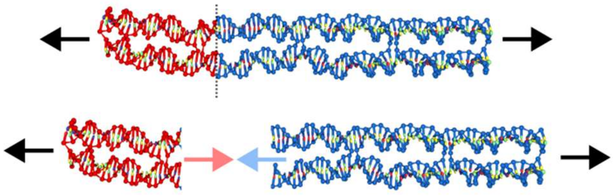

**Figure S9.** Illustrative representation of the methodology to calculate forces in oxDNA. The red and blue colors represent nts upstream and downstream of the section plane.

**S4.5 Verification of force calculation methodology with 2HB DNA**

To verify the force calculation methods, we chose a 2HB rod with a staple layout and length (84 nts) similar to the constituent edges in our metastructure. We applied constant forces (both compressive and tensile) along the axial direction (*z* direction). Separating the edge into 42 sections spaced evenly every 2 nts throughout the structure, we calculated the local stress at each location, by averaging the instantaneous forces over $80\times10^6$ steps. Since there were no forces in the other directions, the forces along *x* and *y* directions are expected to be close to zero. The estimated force should be comparable to the constant magnitude applied, which is given as input to the simulation. The estimates yielded values close to the actual forces applied as shown in Fig S10. Therefore, this method can provide realistic estimates for DNA nanostructures as well.

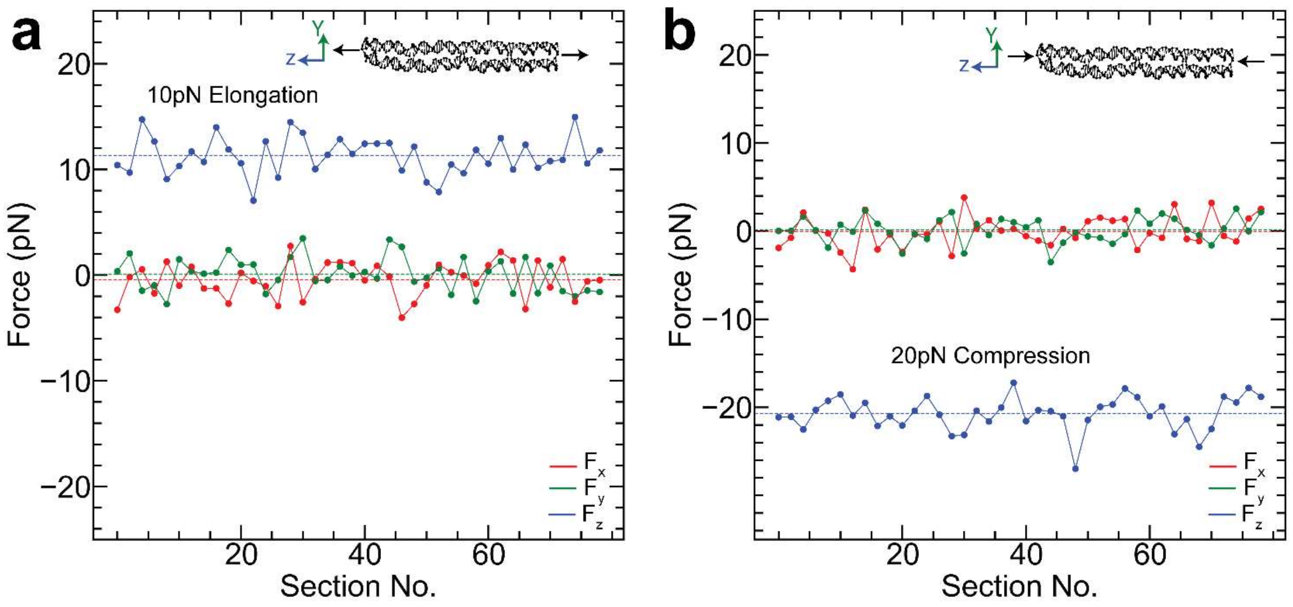


**Figure S10.** Computation of forces on a 2HB edge. (**a**) Forces computed under 10 pN tensile force. The blue line represents the *z* direction while red and green denote *x* and *y* directions, respectively. (**b**) Calculated force estimates when subject to 20 pN compression. In both cases, the calculated average force is in close agreement with the actual force applied.

**S5 Free Energy Calculations**

Traditional MD simulations often fail to explore the whole conformation space (i.e., a wide range of non-equilibrium edge lengths or unfavorable states) due to the high possibility of remaining stuck in equilibrium states. One method of overcoming this shortfall is by applying a bias potential, forcing the structure to explore different conformations by using umbrella sampling. This is achieved by using an order parameter, a structural property that is varied over multiple windows in a series of simulations. For the metastructures in this work, the critical difference is in the mechanical state of edge no. 27 (the edge that buckles in the inadaptable state). We set the end-to-end distance of this edge in both states as the order parameter. We use the ligated structures from the loading simulations as the input for the umbrella sampling simulations. A harmonic biasing force is used

$$V = \frac{k\left(L - L_o^i\right)^2}{2}$$

where $V$ is the bias potential exerted by a spring of stiffness $k$ = 11.4 pN/nm. $L$ and $L_o^i$ are the end-to-end distance and the set value for window $i$ to bias over 80 simulation windows (each run for $25\times10^6$ steps), varying the length of the edge $L_0^i$ from 6.8 to 34 nm. The range of order parameter was chosen such that we scanned beyond the lengths of edges observed during bending (~18 nm) and full extension (~28 nm). The first $5\times10^6$ steps were discarded from all the windows to allow the structure to reach equilibrium. We then used the weighted histogram analysis method (WHAM) to extract the final free energy profile after unbiasing the simulation.

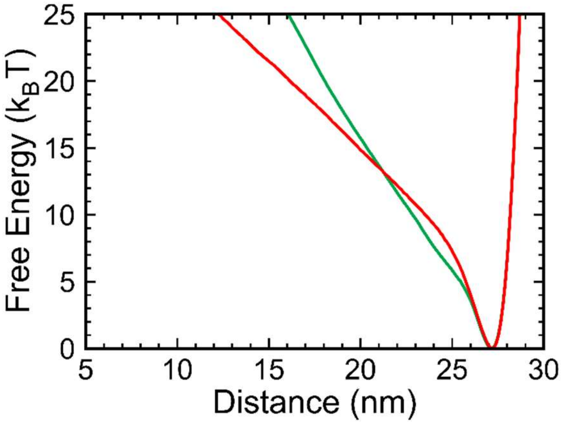


**Figure S11.** Free energy profiles of a 2HB edge (same as that discussed in section S4.5) and the adaptable mode colored red and green, respectively. Both exhibit a single minimum in free energy profiles.

### S6 Bending and Out-of-plane Deformation

The inherent helicity of DNA as well as fluctuations in solution phase often make planar DNA origami by design take up non-planar configurations. However, commonly used imaging techniques such as AFM and transmission electron microscopy (TEM) fail to capture 3D conformations in solution because they require the deposition of the sample on a flat substrate like mica or TEM grids. We accounted for this limitation in our MD simulations by using spring-like repulsive force planes. Using a pair of force planes in conjunction enables the application of a planar 'packing' force, which allows for simulation of the structural configurations comparable to those observed under AFM. All simulations were performed with packing forces using a spring constant of $k = 10^{-3}$ simulation units (about 0.057 pN/nm), unless stated otherwise.

It should be noted that while force planes limit 3D deformations, the mechanical responses of the structures do not change even at lower packing forces. We simulated the loading of DNA metastructures with a minimal packing force. Examining the final states (Fig. S12), the adaptable mode has the conformation consistent with that shown in Fig. 1f, suggesting the packing force has an insignificant impact in the adaptable state. In contrast, the inadaptable mode still exhibits evident buckling. The edge no. 27 clearly protrudes out of the plane in the side and perspective views.

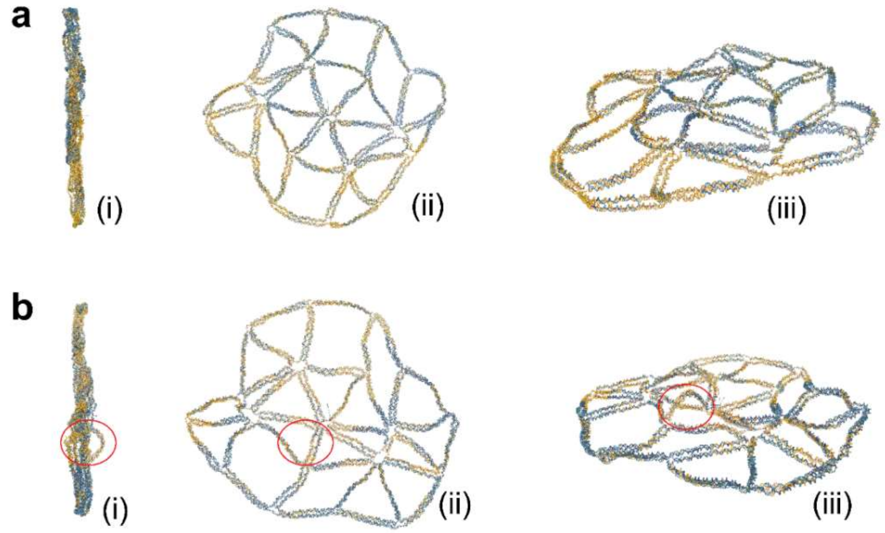


**Figure S12.** Final conformation states obtained from loading simulations with minimal packing force. (**a**) side (i), front (ii) and perspective (iii) views of the adaptable state. (**b**) Corresponding views of the inadaptable state, with one edge sticking out (no. 27 circled in red) experiencing out-of-plane buckling.

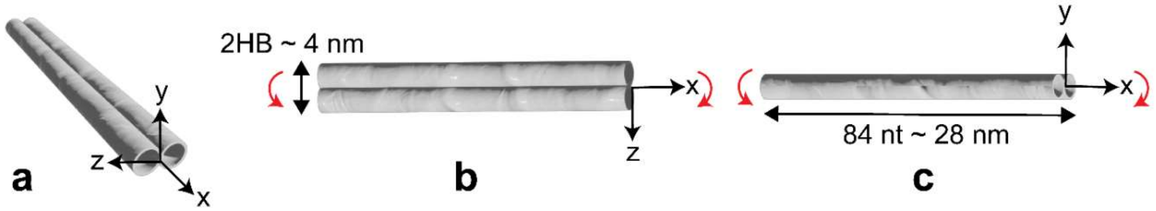


**Figure S13.** (**a**) Different bending modes of a 2HB edge approximated as an elastic beam. The edge can be bent by applying moments along either $y$ (**b**) or $z$ (**c**) axes with a different bending stiffness along each direction.

The deformation behaviors in Fig. S12 can be explained by estimating the bend stiffness of our 2HB edges using elastic beam theory. Assuming the 2HB edge to be a perfectly elastic beam, the geometry of the cross section is not isotropic, with its width (~4 nm) being twice the height of the edge (~2 nm). Therefore, the beam will have distinct bending stiffness values when bent along different directions (Fig. S13). The edge can be bent by applying moment along either the *y* or *z* axes which correspond to in-plane and out of plane deformations in the context of our structure. Prior studies have estimated that the Young's modulus of a DNA duplex is around 0.3 GPa.[11] For the calculations based on the parallel axis theorem, we approximate the diameter (*d*) of a DNA double helix to be 2 nm and the Young's modulus (*E*) to be 276 MPa (or 276 pN/nm$^2$). We use the bending stiffness $\mathrm{EI}$ to compare both cases, where $\mathrm{I}$ the area moment of inertia.

For the beam in Fig. S13b, the bending moment along the *y* direction is:

$$I_{yy} = 2 \cdot \frac{\pi d^4}{64} + 2 \cdot \frac{\pi d^4}{16} = 7.85\, nm^4$$
$$EI = 2167\, pN \cdot nm^2$$

For the bending moment along the *z* direction (Fig. S13c),

$$I_{zz} = 2 \cdot \frac{\pi d^4}{64} = 1.57\, nm^4$$
$$EI = 434\, pN.nm^2$$

Clearly, the stiffness of the edge when bent in-plane with the moment along the *y* axis is far greater than that with the moment along the *z*-axis (bending out-of-plane). This discrepancy in stiffness makes it feasible for the edge to buckle in the latter direction. This phenomenon is consistent with the packing force of $k = 10^{-3}$, where the cross section of edge 27 rotates and still buckles along the most favored direction in 2D plane as shown in Figs. 1 and 3.

### S7 Additional AFM Images

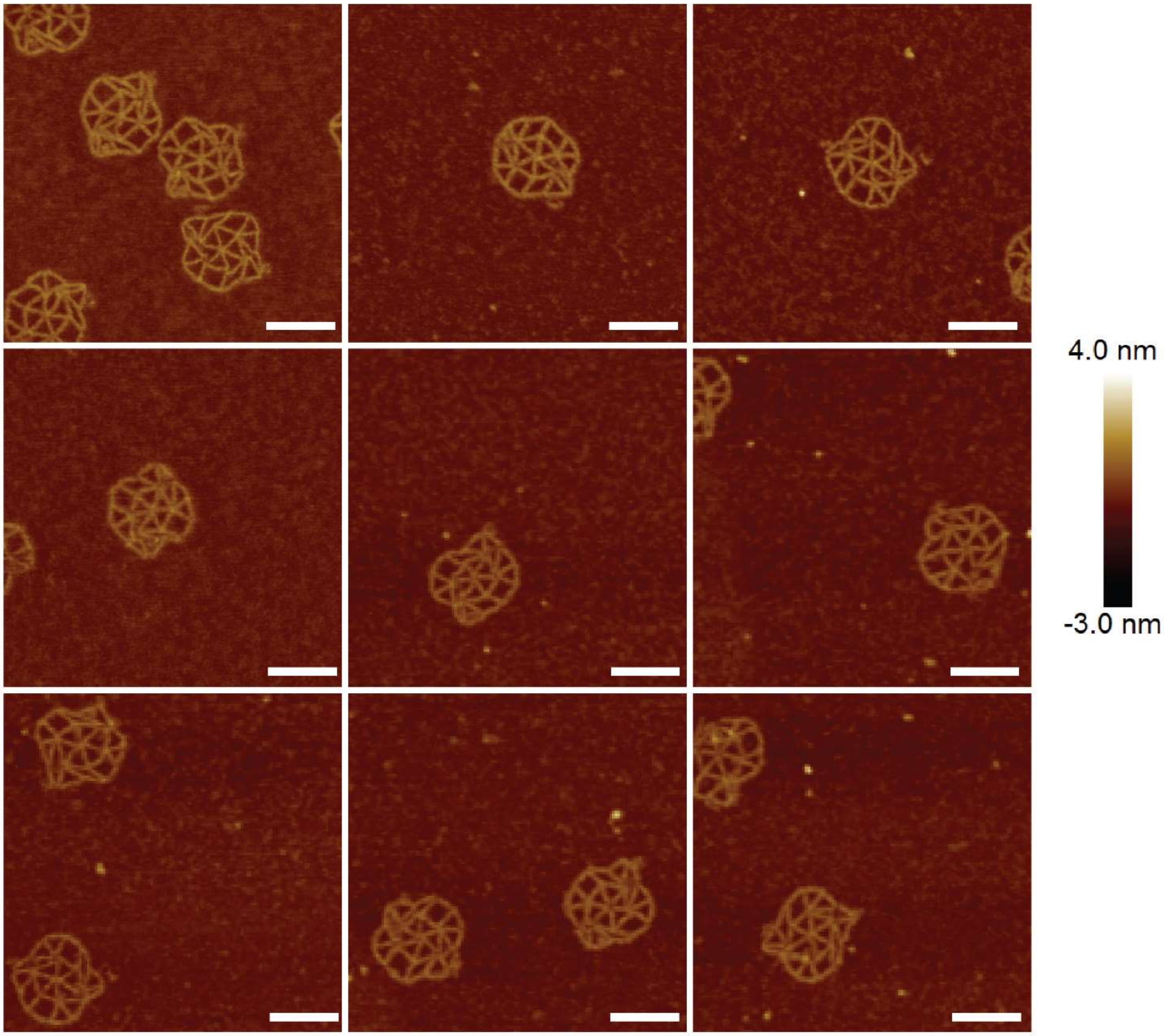


**Figure S14.** AFM images of DNA origami in the initial (undefined) state with all jack edges absent corresponding to Fig. 2a (i). Scale bar: 100 nm.

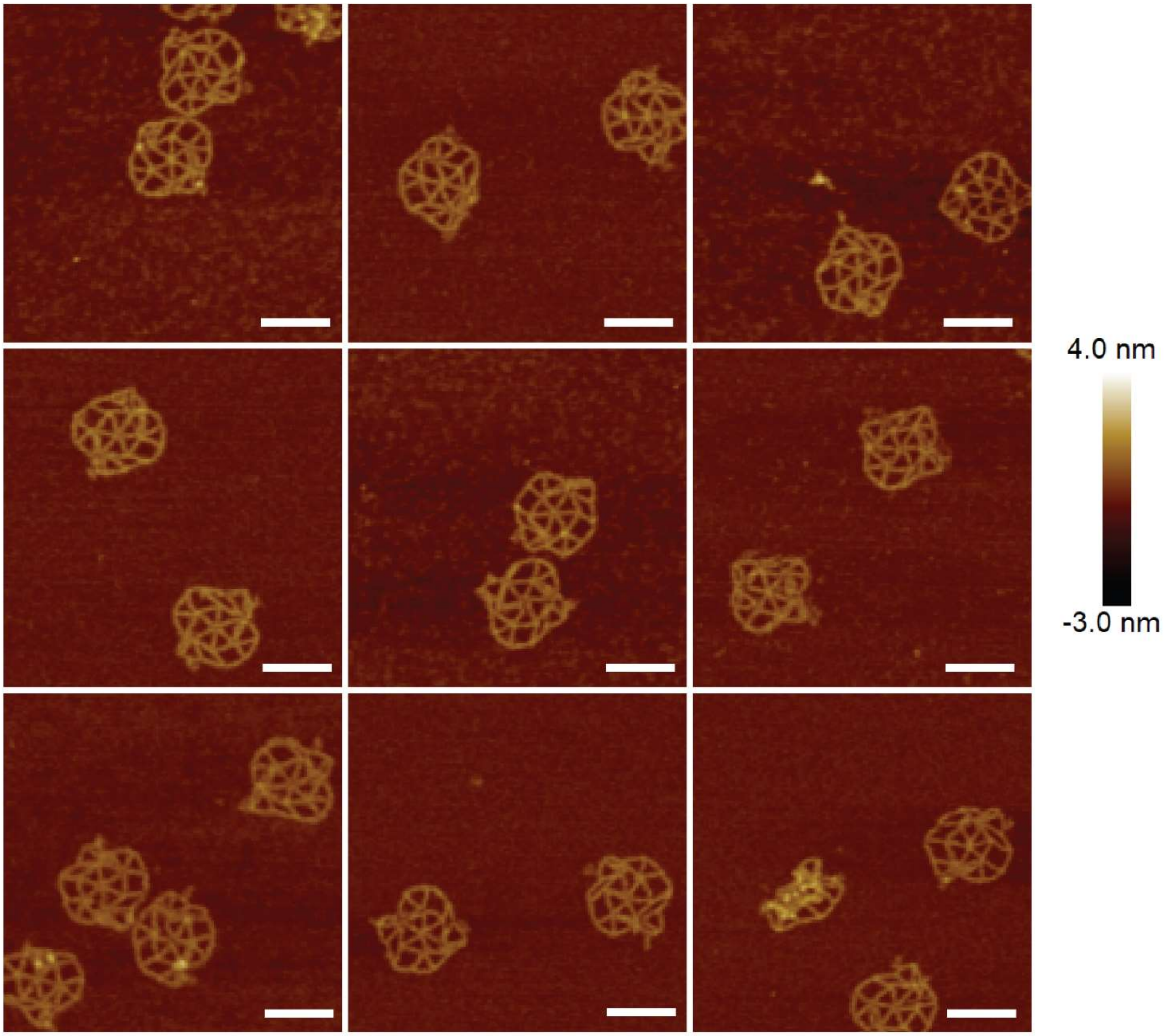


**Figure S15.** AFM images of adaptable DNA metastructures which correspond to Figs. 2a (ii) and 2b. Scale bar: 100 nm.

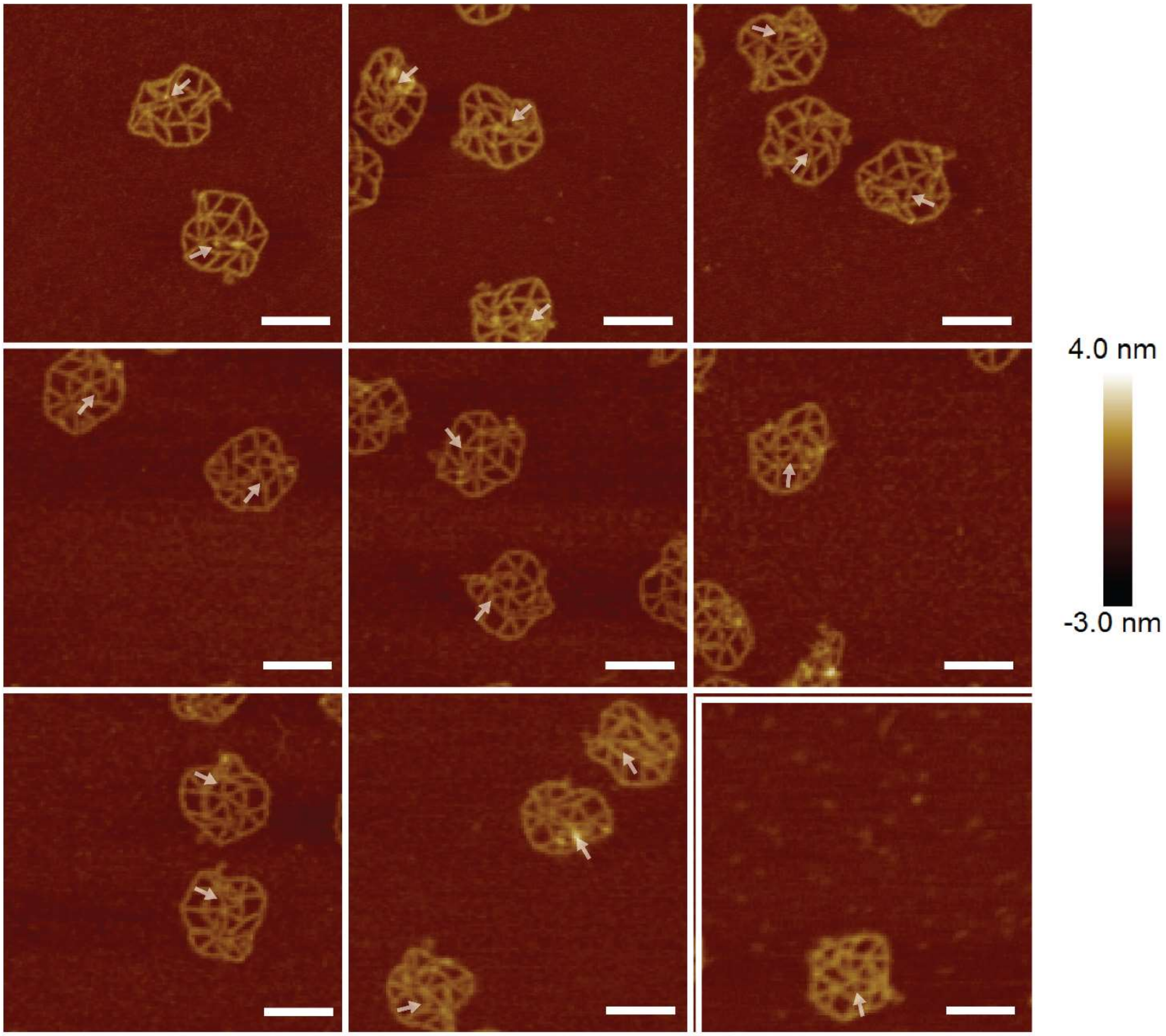


**Figure S16.** AFM scans of inadaptable DNA metastructures. These are additional images to those in Figs. 2a (iii) and 2c. White arrows indicate the buckled edges. Scale bar: 100 nm.

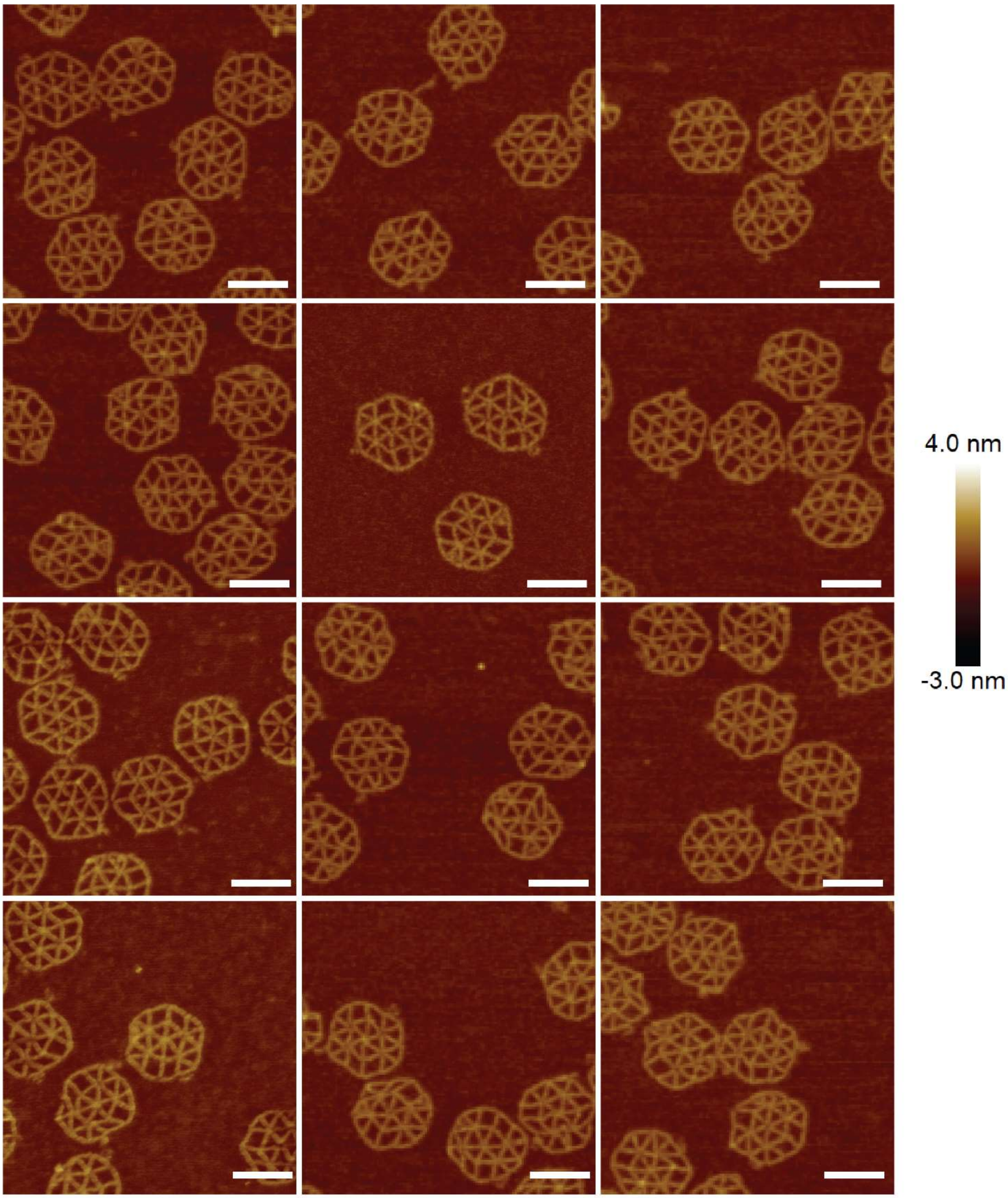


**Figure S17.** AFM images of DNA metastructures in the initial (extended) state with all jacks extended. These images are consistent with those in Figs. 2e (i) and 2f. Scale bar: 100 nm.

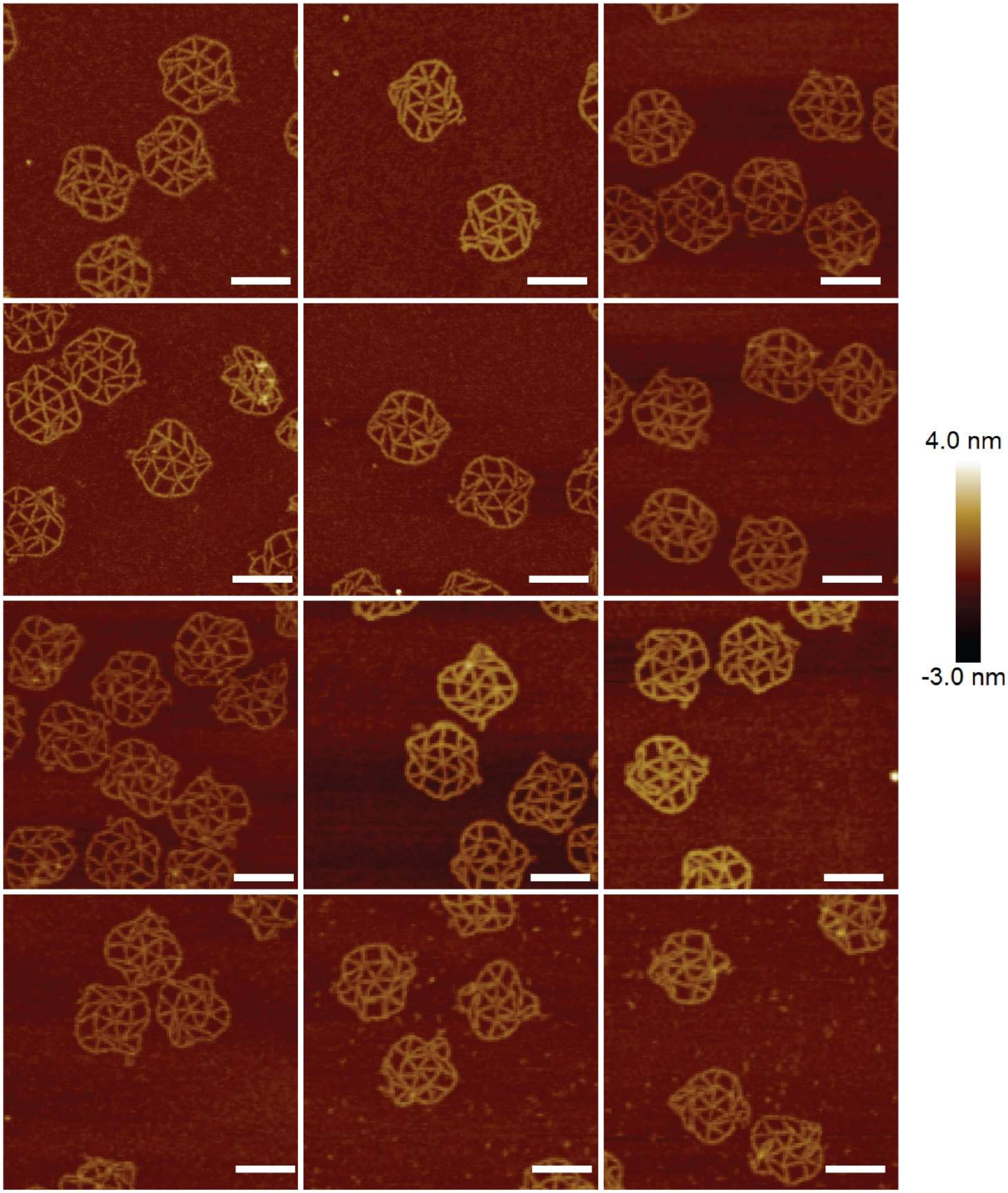


**Figure S18.** AFM images of the adaptable undefined state (Fig. 2e (ii)), where adaptable jack edges are removed by using jack releasers from the initial extended state (Fig. 2e (i)). Scale bar: 100 nm.

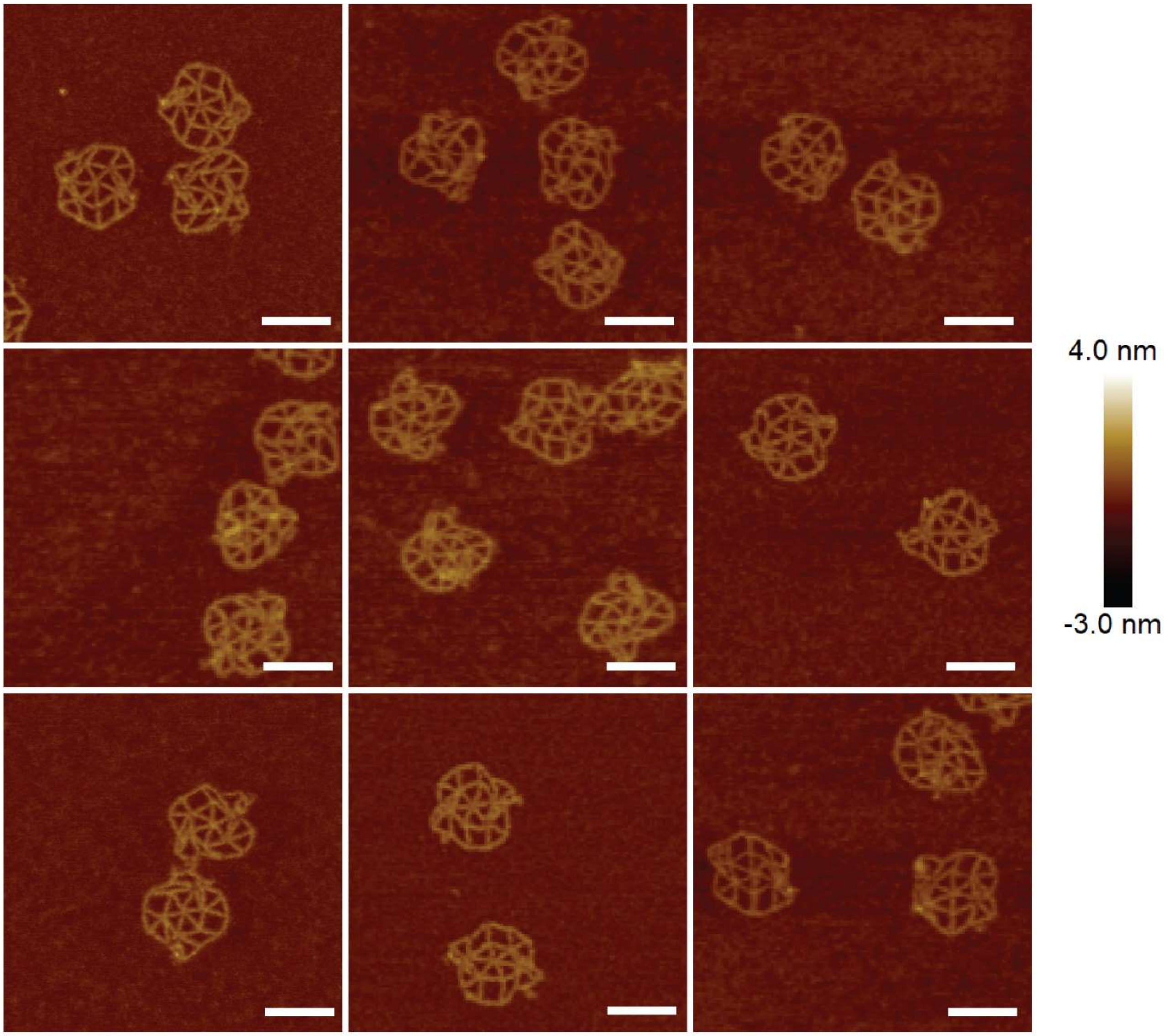


**Figure S19.** AFM scans of adaptable DNA metastructures which are consistent with Figs. 2e (iii) and 2g. Scale bar: 100 nm.

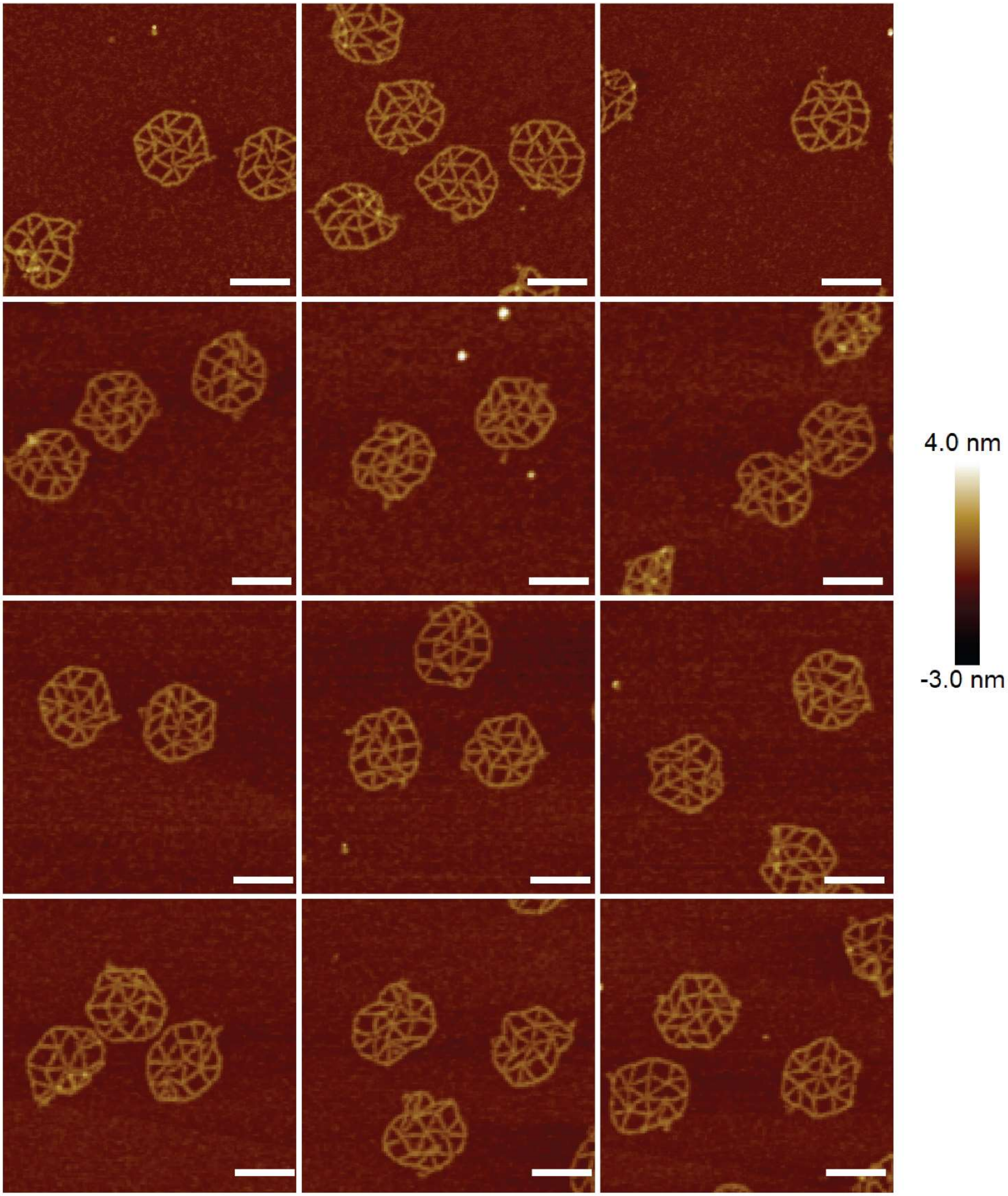


**Figure S20.** AFM scans of the inadaptable undefined state (Fig. 2e (iv)), where inadaptable jack edges are removed by introducing inadaptable jack releasers. Scale bar: 100 nm.

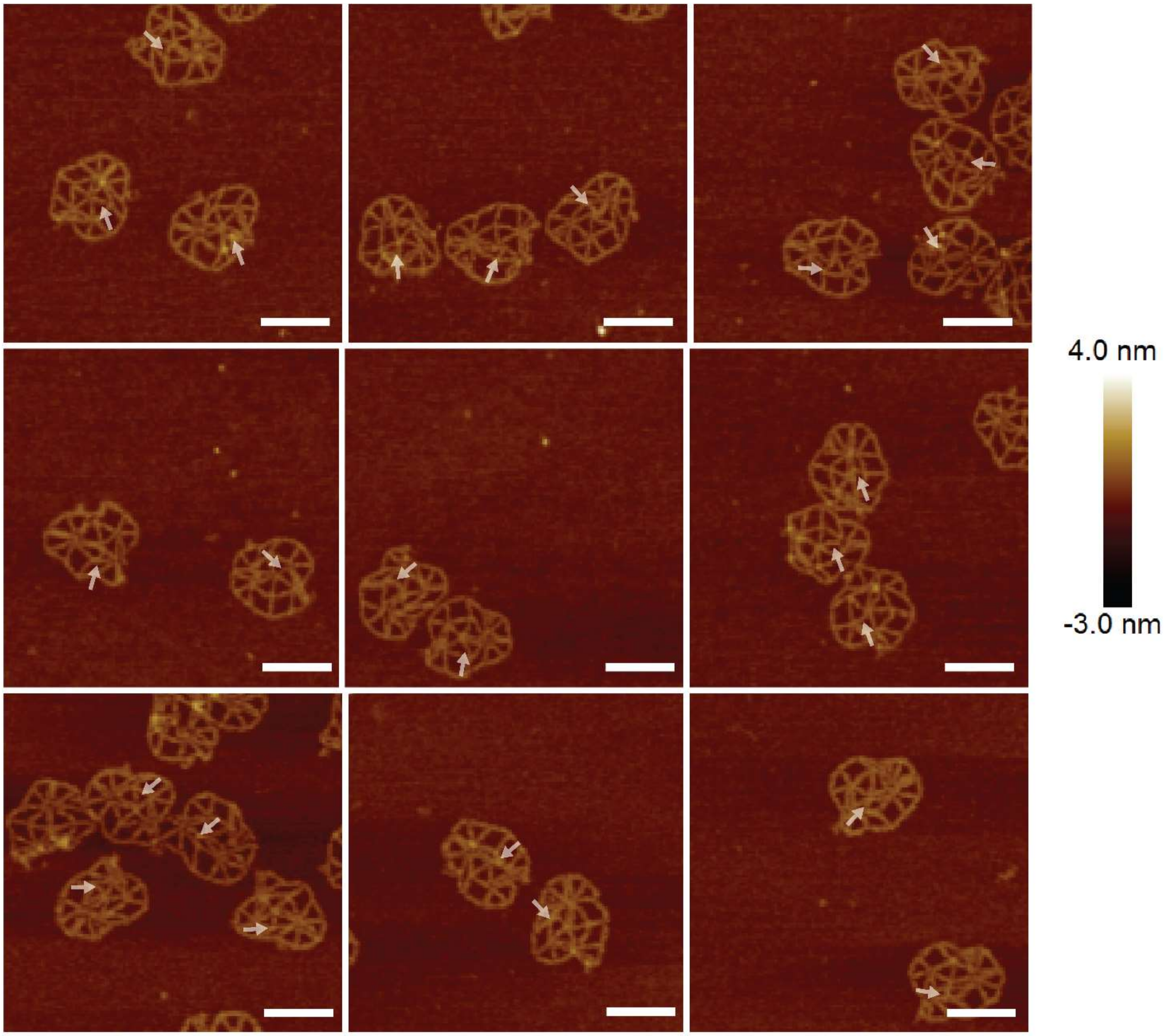


**Figure S21.** AFM images of inadaptable DNA metastructures which correspond to Figs. 2e (v) and 2h. White arrows indicate the buckled edges. Scale bar: 100 nm.

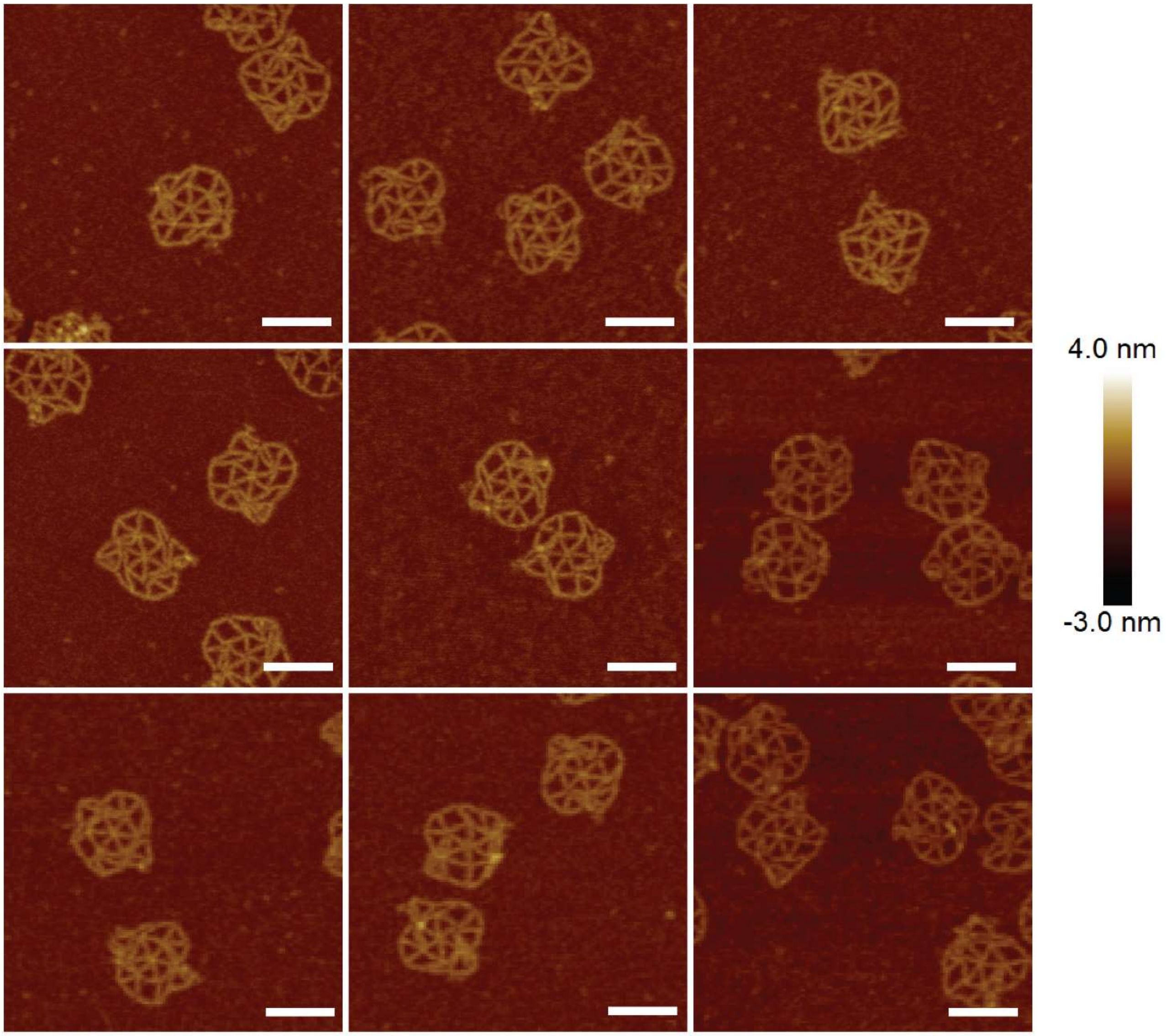


**Figure S22.** AFM scans of adaptable DNA metastructures reconfigured from the inadaptable state (Fig. 2e (v) → (iii)). Scale bar: 100 nm.

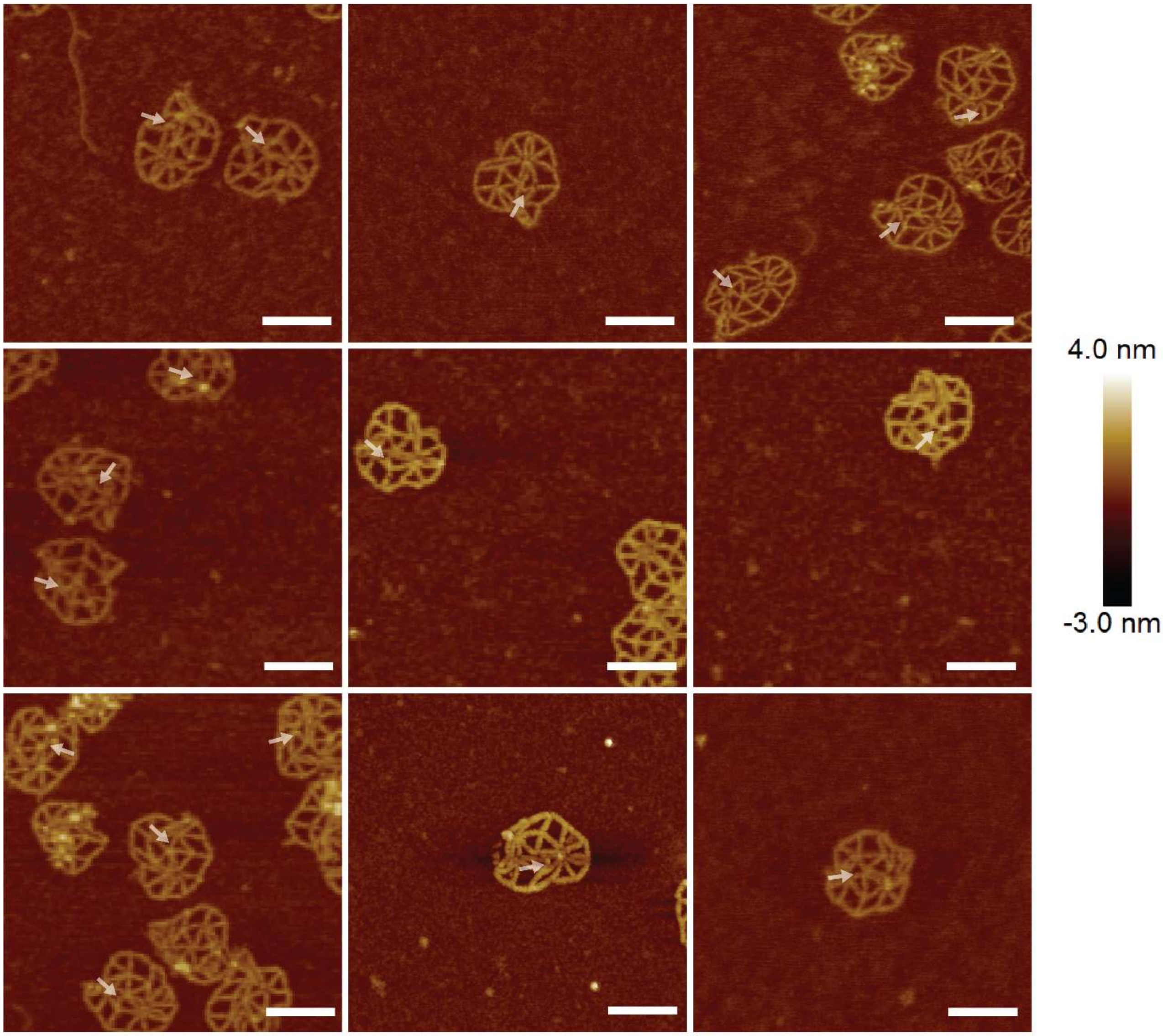


**Figure S23.** AFM images of inadaptable DNA metastructures after switching from the adaptable state (Fig. 2e (iii) → (v)). White arrows indicate the buckled edges. Scale bar: 100 nm.

**S8 DNA Sequences**

| Regular Edge Staples | |
|---|---|
| Name | Sequence |
| S[1][1] | GTTAATGCCCTACATGGCTTTTGATGATACACCGTATAAACA |
| S[1][2] | AACAGTGCGGAGTGTACTGGTAATAAGTTTTAACAGAGCCAC |
| S[1][3] | CACCCTCACAGAACCGCCACCCTCGGGGTCAGTGCCTTGAGT |
| S[1][4] | TAGCAAGCCCACACCCTCAGAACCGCCACCCTTTTTCAGGGA |
| S[2][1] | GATCTAAAGTATGTACCGTAACACTGAGTTTGTTAGCGTAAC |
| S[2][2] | CCCTCATACGTCACCAGTACAAACTACAACGCCTAAGGCCGC |
| S[2][3] | TTTTGCGGGGCTTGCAGGGAGTTAGTAGCATTCCACAGACAG |
| S[2][4] | CTCAGCAGCGAACCGATATATTCGGTCGCTGAGATCGTCACC |
| S[3][1] | GCCCCCTTATGCAGCACCGTAATCAGTAGCGTTTCGGTCATA |
| S[3][2] | ATCGGCATACAGAATCAAGTTTGCCTTTAGCGTCGCCAGAAT |
| S[3][3] | GGAAAGCGATAAATCCTCATTAAAAGACTGTAGCGCGTTTTC |
| S[3][4] | ATTTACCGTTCGGCCTTGATATTCACAAACAACAGTCTCTGA |
| S[4][1] | GCAGGTCAGACTATTTCGGAACCTATTATTCAGGAGGTTGAG |
| S[4][2] | AAGTATTAAGAAGAGCCGCCGCCAGCATTGACTGAAACATGA |
| S[4][3] | AACCACCACCGGCTGAGACTCCTCAAGAGAAGCCGCCACCAG |
| S[4][4] | TAGCGGGGTTTCTCAGAGCCACCACCCTCAGAGGATTAGGAT |
| S[5][1] | AGTGAGAATAAGTACCAGGCGGATAAGTGCCAGTTTCAGCGG |
| S[5][2] | TTGATATAAGTTTTTGCTAAACAACTTTCAACGTCGAGAGGG |
| S[5][3] | CTGTATGGGAATAGCCCGGAATAGGTGTATCAAATGAATTTT |
| S[5][4] | GGAGGTTTAGTGTCGTCTTTCCAGACGTTAGTACCGTACTCA |
| S[6][1] | CGCGAAACAAGTTTATCAGCTTGCTTTCGAGATTATACCAAG |
| S[6][2] | TAAACAGCTTGACTCATCTTTGACCCCCAGCGGTGAATTTCT |
| S[6][3] | ACACTAAAACATACCGATAGTTGCGCCGACAGGCAAAAGAAT |
| S[6][4] | CCATCGCCCACGCACCAACCTAAAACGAAAGAATGACAACAA |
| S[7][1] | AATGCCACTAGAACGAGGGTAGCAACGGCTAGTAAAATACGT |
| S[7][2] | TTAAACGGCAGAGGCTTTGAGGACTAAAGACTTTCAAATCAA |
| S[7][3] | CGTAACAACCGGATATTCATTACCTTCATGAGGAAGTTTCCA |
| S[7][4] | TCAGTGAATAACAAGAGTAATCTTGACAAGAAAGCTGCTCAT |
| S[8][1] | GTCTGAGAGATTTGCCATCTTTTCATAATCACAAAATCATAG |
| S[8][2] | AACCAGAGCCAAAGAGTCAATAGTGAATTTATAAATCACCGG |

| | |
|---|---|
| S[8][3] | AGACGCTGAGCCACCGGAACCGCCTCCCTCAAGCTTAGATTA |
| S[8][4] | CCTCAGAACCGAATCCTTGAAAACATAGCGATGAGCCGCCAC |
| S[9][1] | GCAGAGGCGAGAACAACTAAAGGAATTGCGATACAAAATCGC |
| S[9][2] | TTTCACGTTGATGATTGCTTTGAATACCAAGTATAATAATTT |
| S[9][3] | CGGATTCGCCAAATCTCCAAAAAAAAGGCTCAGAAACAATAA |
| S[9][4] | CTTTAATTGTAAACAGTACCTTTTACATCGGGCAAAAGGAGC |
| S[10][1] | CAATTTTATCCGCGAGGCGTTTTAGCGAACCGCACCCAGCTA |
| S[10][2] | GCTATTTTTCCCGACTTGCGGGAGGTTTTGAAGCCATTACCA |
| S[10][3] | TTAGCAAGAAATCACCAGTAGCACCTTAAATCAAGATTAGTT |
| S[10][4] | TCACCAATGAAATTTGGGAATTAGAGCCAGCAGCCGGAAACG |
| S[13][1] | TTACCTTTTTTCATTTCAATTACCTGAGCAATTTCATTTGAA |
| S[13][2] | TTTAACAAAAGAAGATGATGAAACAAACATCAAGGAAAGGAA |
| S[13][3] | TTGAGGAATGGCAAATCAACAGTTAAAACAAAATTAATTACA |
| S[13][4] | AATATCTTTAGCTCAATCAATATCTGGTCAGTGGTTATCTAA |
| S[14][1] | CGTATTAAATCTAACAACTAATAGATTAGAGATTCGACAACT |
| S[14][2] | TACAAACACCGTCAATAGATAATACATTTGAGGAATTGCGTA |
| S[14][3] | GATTTTCAAAACAGAAATAAAGAATTTAGAAGTATTAGACTT |
| S[14][4] | CAGATGAATATTATCAAAATTATTTGCACGTAGGTTTAACGT |
| S[15][1] | GGTTAGAACCACAACGGAGATTTGTATCATCAATAATGGAAG |
| S[15][2] | TTGTGTCGAAAATTGTTTGGATTATACTTCTGGCCTGATAAA |
| S[15][3] | TATAATCCTGTCCGCGACCTGCTCCATGTTACAATTCATCAA |
| S[15][4] | ACGAGGCGCAGTTCCTGATTATCAGATGATGGCTTAGCCGGA |
| S[16][1] | TAGAAAGATTCAATCATAAGGGAACCGAACTATTATTACAGG |
| S[16][2] | GAAAGAGGACAAAAACGAACTAACGGAACAACGACCAACTTT |
| S[16][3] | CTACGTTAATGATGAACGGTGTACAGACCAGGGAAGAAAAAT |
| S[16][4] | TGGCTGACCTTATTATACCAGTCAGGACGTTGGCGCATAGGC |
| S[17][1] | TTTAAGAACTGACGAGAAACACCAGAACGAGCCTTATGCGAT |
| S[17][2] | GTGAATTATAGTAAATTGGGCTTGAGATGGTTTATTAGATAC |
| S[17][3] | ATTTCGCAAGATTTAGTTTGACCAATTTCAACTTTAATCATT |
| S[17][4] | AACCTGTTTAGTTCCCAATTCTGCGAACGAGTAATGGTCAAT |
| S[18][1] | TAAGAAACGATCTTACCAACGCTAACGAGCGCCCAATCCAAA |
| S[18][2] | TTATTTATTCTTTCCAGAGCCTAATTTGCCAGTTCAATAGAA |
| S[18][3] | AATTCATATTTATTTTGTCACAATACAAAATAAACAGCCATA |

| | |
|---|---|
| S[18][4] | GCGCCAAAGACGCAAAGACACCACGGAATAAGTGGTTTACCA |
| S[19][1] | AGGGCTTAATGGAATCATAATTACTAGAAAACGCTCAACAGT |
| S[19][2] | AAAGCCAAAGCCTGTTTAGTATCATATGCGTTATAAAATCTA |
| S[19][3] | AAGCATCAAGCCAGCAGCAAATGAACAAATTCTTACCAGTAT |
| S[19][4] | CCTCAAATATCCTGCAACAGTGCCACGCTGAGCCTTGCTGAA |
| S[20][1] | GGGACATTCTGCCCGAACGTTATTAATTTTACAGTAATAAAA |
| S[20][2] | TAACATTATCACAGATTCACCAGTCACACGACAAAGTTTGAG |
| S[20][3] | TTTACATTGGTTTTGCGGAACAAAGAAACCAGAAATGGATTA |
| S[20][4] | CGGAATTATCACATTTTGACGCTCAATCGTCTCCAGAAGGAG |
| S[21][1] | CATAACCCTCGTTGAGATTTAGGAATACCACAGCAACACTAT |
| S[21][2] | ATAGTAAGATTCAACTAATGCAGATACATAACGCTAAAGTAC |
| S[21][3] | GGTGTCTGGTTTTAAATATGCAACCAAAAGGAATTACGAGGC |
| S[21][4] | TCCATATAACATATAATGCTGTAGCTCAACATGAAGTTTCAT |
| S[22][1] | GGCTTATCCGTGTTTAACGTCAAAAATGAAACAGATATAGAA |
| S[22][2] | TTACAGAGAGACGCGCCCAATAGCAAGCAAATATAGCAGCCT |
| S[22][3] | GAATCATTACATAACATAAAAAAACCGAGGATTTCATCGTAG |
| S[22][4] | TAACGGAATACGAACAAGCAAGCCGTTTTTATAACGCAATAA |
| S[23][1] | GAACAAGAAAAGAACTGGCATGATTAAGACTAGATAAGTCCT |
| S[23][2] | CAGTATGTTAGACGCGCCTGTTTATCAACAATCCTTATTACG |
| S[23][3] | CTAATGCAGACAAACGTAGAAAATACATACAAACATGTTCAG |
| S[23][4] | AACATATAAAAGTCCAGACGACGACAATAAACTAAAGGTGGC |
| S[24][1] | AGGTAAAGTAATCGCCATATTTAACAACGCCGTACCGACAAA |
| S[24][2] | AATATAAAAACATGTAATTTAGGCAGAGGCATTTAGGGCGCT |
| S[24][3] | GGCAAGTGGAAAGGAGCGGGCGCTTCGAGCCAGTAATAAGAG |
| S[24][4] | GCTGCGCGTAAGAGAAAGGAAGGGAAGAAAGCTAGCGGTCAC |
| S[25][1] | ATCAGAGCGGACACCCGCCGCGCTTAATGCGTCCTCGTTAGA |
| S[25][2] | ACGTGCTTCCGCTACAGGGCGCGTACTATGGTTGCAGAAGAT |
| S[25][3] | AAAACAGACCGAACGAACCACCAGCTTTGACGAGCACGTATA |
| S[25][4] | TCAGTATTAACAAAACATCGCCATTAAAAATAGGTGAGGCGG |
| S[26][1] | CGAACTGATAACAGAGATAGAACCCTTCTGATCTTTAATGCG |
| S[26][2] | GCTATTAGCCTGAAAGCGTAAGAATACGTGGCACAATACTTC |
| S[26][3] | TTTGATTAAATTAACCGTTGTAGCAGACAATATTTTTGAATG |
| S[26][4] | CACTTGCCTGAAAAGAGTCTGTCCATCACGCAGTAATAACAT |

| | |
|---|---|
| S[27][1] | ATCGATGAACAGAACTCAAACTATCGGCCTTCAAACAAGAGA |
| S[27][2] | TCCAGAACAATTCATTGCCTGAGAGTCTGGAGGCTGGTAATA |
| S[27][3] | GGCTATCAGGATTACCGCCAGCCATTGCAACAGATCTACAAA |
| S[27][4] | CTCATGGAAATGAGAGGGTAGCTATTTTTGAGAGGAAAAACG |
| S[29][1] | CAAGGCAAAGGCGTCCAATACTGCGGAATCGAATCATACAGG |
| S[29][2] | TCATTGAATCCTAGTAGCATTAACATCCAATATCATAAATAT |
| S[29][3] | CTACTAATAGCCCTCAAATGCTCATTTTTGCTGGCATCAATT |
| S[29][4] | GAGCTTAATTGTTGGGGCGCGAGCTGAAAAGGGGATGGCTTA |
| S[30][1] | TGAGACGGGCAAACAGGAGGCCGATTAAAGGCTTTTCACCAG |
| S[30][2] | AGGAACGGTACTTGGGCGCCAGGGTGGTTTTTGATTTTAGAC |
| S[30][3] | GGTTTGCGTAGCCAGAATCCTGAGAAGTGTTCGGGGAGAGGC |
| S[30][4] | GTGAGGCCACCATTAATGAATCGGCCAACGCGTTTATAATCA |
| S[31][1] | TTAAATTGTATCGTAAAACTAGCATGTCAATTAAGCAAATAT |
| S[31][2] | AGATTGTACATATGTACCCCGGTTGATAATCAGAAACCCTCA |
| S[31][3] | TATATTTTGGATAAAAATTTTTAGAAAGCCCCAAAAACAGGA |
| S[31][4] | CCTGAGTAATGGAAGCCTTTATTTCAACGCAAAAATGCAATG |
| S[32][1] | CCAAGTACCGAATATCCCATCCTAATTTACGCGGGTATTAAA |
| S[32][2] | TCCAAGAAAGCATGTAGAAACCAATCAATAATCGCGATGGCC |
| S[32][3] | CACTACGTAAACCGTCTATCAGGGGCTGTCTTTCCTTATCAT |
| S[32][4] | CCAAATCAAGTACTCCAACGTCAAAGGGCGAAGAACCATCAC |
| S[33][1] | AAAATCCTGTGGGGTCGAGGTGCCGTAAAGCCCCAGCAGGCG |
| S[33][2] | AACCCTAAAGGAAGCGGTCCACGCTGGTTTGCACTAAATCGG |
| S[33][3] | GAGTTGCAGCGAGCCCCCGATTTAGAGCTTGTGGCCCTGAGA |
| S[33][4] | CCGGCGAACGTAGCTGATTGCCCTTCACCGCCACGGGGAAAG |
| S[34][1] | AACGCCAGGGCTGGGGTGCCTAATGAGTGAGTTAAGTTGGGT |
| S[34][2] | TTAATTGCGTTGGGGGATGTGCTGCAAGGCGACTAACTCACA |
| S[34][3] | GCTGGCGAAAGCGCTCACTGCCCGCTTTCCACTATTACGCCA |
| S[34][4] | CTGTCGTGCCACGATCGGTGCGGGCCTCTTCGGTCGGGAAAC |
| S[35][1] | AACTGTTGGGTAATATTTTGTTAAAATTCGCTCAGGCTGCGC |
| S[35][2] | TGTTAAATCAGAGGCAAAGCGCCATTCGCCATATTAAATTTT |
| S[35][3] | GCCGGAAACCCTCATTTTTTAACCAATAGGACCGCTTCTGGT |
| S[35][4] | AAATAATTCGCACTCCAGCCAGCTTTCCGGCAACGCCATCAA |
| S[36][1] | GACCGTAATGGCCTTCCTGTAGCCAGCTTTCAACGGCGGATT |

| | |
|---|---|
| S[36][2] | TGGGAACAATCAACATTAAATGTGAGCGAGTAACGTTGTACC |
| S[36][3] | AAAAACATGCATAAAGCTAAATCGAACCCGTCGGATTCTCCG |
| S[36][4] | TAATACTTTTGATTAAGCAATAAAGCCTCAGATATGACCCTG |
| S[37][1] | AGTCCACTATGGTGGTTCCGAAATCGGCAAATTTGGAACAAG |
| S[37][2] | TGTTCCAGATCCCTTATAAATCAAAAGAATAGCCTTCCACAC |
| S[37][3] | AACATACGTGTTATCCGCTCACAACGAGATAGGGTTGAGTGT |
| S[37][4] | ATAAAGTGTAATAGCTGTTTCCTGTGTGAAATAGCCGGAAGC |
| S[38][1] | CGAATTCGTACCAGTCACGACGTTGTAAAACGGTACCGAGCT |
| S[38][2] | GATCCCCGGACGGCCAGTGCCAAGCTTGCATGCCAGTTTGAG |
| S[38][3] | GGGACGACTAACCGTGCATCTGCCTGCAGGTCGACTCTAGAG |
| S[38][4] | GCCTCAGGAAGTTGGTGTAGATGGGCGCATCGGACAGTATCG |

| Extended Jack Staples | |
|---|---|
| Name | Sequence |
| JS[11][1] | GTCACCGACTTTTTTTAACCTCCTCGCAGC |
| JS[11][2] | AATTATCACCCGGCTTAGGTTGGGTTATATACATTAAAGGTGCTATTAAA |
| JS[11][3] | AATGCTGATGCTAAATATTGACGGAAATTATTACTATATGTACCGACTTG |
| JS[11][4] | GGAGGGAAGGAAATCCAATCGCAAGACAAAGAACCGATTGAGGCTGCACC |
| JS[11][5] | AACGCGAGAAGGCGACATTCGCGGTACA |
| JS[12][1] | AATAAGAATAATTTCAAATATAATCAGCAT |
| JS[12][2] | TAAGGCGTTATTTTAGTTAATTTCATCTTCTCGTGTGATAAACTGCGAAG |
| JS[12][3] | ATACCGACGACCTAAATTTAATGCTTGCTTCTGTAAATCGTCACTAAGCT |
| JS[12][4] | GCTATTAAATAAATCAATATATGTGAGTGAATAACGTTTGAACCATCCAG |
| JS[12][5] | TTAATTTTCCGAAACAGTACACTCAGCC |
| JS[28][1] | TGATAAATTAACCAGACGACGATTAAAACT |
| JS[28][2] | CCGTTCTAGCTAAAAACCAAAATAGCGAGAGATGATATTCAACACTGAAG |
| JS[28][3] | AAGAAGTTTTGGACAGTCAAATCACCATCAATGCTTTTGCAATAACAGCC |
| JS[28][4] | AAAGGCCGGACCAGAGGGGGTAATAGTAAAAAAAAGGGTGAGTGGATAGC |
| JS[28][5] | TGTTTAGACTGTAAAGATTCCTCCGAGT |

| Extended Jack Releasers | |
|---|---|
| Name | Sequence |
| RJS[11][1] | GCTGCGAGGAGGTTAAAAAAAGTCGGTGAC |
| RJS[11][2] | TTTAATAGCACCTTTAATGTATATAACCCAACCTAAGCCGGGTGATAATT |
| RJS[11][3] | CAAGTCGGTACATATAGTAATAATTTCCGTCAATATTTAGCATCAGCATT |
| RJS[11][4] | GGTGCAGCCTCAATCGGTTCTTTGTCTTGCGATTGGATTTCCTTCCCTCC |
| RJS[11][5] | TGTACCGCGAATGTCGCCTTCTCGCGTT |
| RJS[12][1] | ATGCTGATTATATTTGAAATTATTCTTATT |
| RJS[12][2] | CTTCGCAGTTTATCACACGAGAAGATGAAATTAACTAAAATAACGCCTTA |
| RJS[12][3] | AGCTTAGTGACGATTTACAGAAGCAAGCATTAAATTTAGGTCGTCGGTAT |
| RJS[12][4] | CTGGATGGTTCAAACGTTATTCACTCACATATATTGATTTATTTAATAGC |
| RJS[12][5] | GGCTGAGTGTACTGTTTCGGAAAATTAA |
| RJS[28][1] | AGTTTTAATCGTCGTCTGGTTAATTTATCA |
| RJS[28][2] | CTTCAGTGTTGAATATCATCTCTCGCTATTTTGGTTTTTAGCTAGAACGG |
| RJS[28][3] | GGCTGTTATTGCAAAAGCATTGATGGTGATTTGACTGTCCAAAACTTCTT |
| RJS[28][4] | GCTATCCACTCACCCTTTTTTTTACTATTACCCCCTCTGGTCCGGCCTTT |
| RJS[28][5] | ACTCGGAGGAATCTTTACAGTCTAAACA |

| Short Jack Staples | |
|---|---|
| Name | Sequence |
| SJ11[1][1] | AATTATCACCGTCACCGACTTGGCGACATTCAACCGATTGAGGATAATTT |
| SJ11[1][2] | CATTAAAGGTGGGAGGGAAGGTAAAACTTT |
| SJ11[2][1] | GCAAGACAAAGAACGCGAGAATTTTTAACCTCCGGCTTAGGTGGCTGAAC |
| SJ11[2][2] | CAAATCCAATCTGGGTTATATACCGTTACG |
| SJ12[1][1] | CGTGTGATAAATATGTGAGTGACCTACGCG |
| SJ12[1][2] | TAAGGCGTTAAATAAGAATAAGAAACAGTACATAAATCAATACGACGTGG |
| SJ12[2][1] | TCTGTAAATCTTTCATCTTCTCGGACAG |
| SJ12[2][2] | GTCGCTATTAATTAATTTTCCTTTCAAATATATTTTAGTTAAGACGCTTC |
| SJ28[1][1] | CCGTTCTAGCTGATAAATTAAGTAAAGATTCAAAAGGGTGAGTGATTCGA |
| SJ28[1][2] | ATGATATTCAAAAAGGCCGGAGAAATGCTC |
| SJ28[2][1] | TAATAGTAAAATGTTTAGACTCCAGACGACGATAAAAACCAAGGTAGGTC |
| SJ28[2][2] | GCCAGAGGGGGAATAGCGAGAGCACCATGC |

| Short Jack Releasers | |
|---|---|
| Name | Sequence |
| RSJ11[1][1] | AAATTATCCTCAATCGGTTGAATGTCGCCAAGTCGGTGACGGTGATAATT |
| RSJ11[1][2] | AAAGTTTTACCTTCCCTCCCACCTTTAATG |
| RSJ11[2][1] | GTTCAGCCACCTAAGCCGGAGGTTAAAAATTCTCGCGTTCTTTGTCTTGC |
| RSJ11[2][2] | CGTAACGGTATATAACCCAGATTGGATTTG |
| RSJ12[1][1] | CGCGTAGGTCACTCACATATTTATCACACG |
| RSJ12[1][2] | CCACGTCGTATTGATTTATGTACTGTTTCTTATTCTTATTTAACGCCTTA |
| RSJ12[2][1] | CTGTCCGAGAAGATGAAAGATTTACAGA |
| RSJ12[2][2] | GAAGCGTCTTAACTAAAATATATTTGAAAGGAAAATTAATTAATAGCGAC |
| RSJ28[1][1] | TCGAATCACTCACCCTTTTGAATCTTTACTTAATTTATCAGCTAGAACGG |
| RSJ28[1][2] | GAGCATTTCTCCGGCCTTTTTGAATATCAT |
| RSJ28[2][2] | GCATGGTGCTCTCGCTATTCCCCCTCTGGC |
| RSJ28[2][1] | GACCTACCTTGGTTTTTATCGTCGTCTGGAGTCTAAACATTTTACTATTA |

### S9 References


1. Meeussen, A.S., Oğuz, E.C., Shokef, Y. & Hecke, M.v. Topological Defects Produce Exotic Mechanics in Complex Metamaterials. *Nature Physics* **16**, 307-311 (2020).
2. Douglas, S.M. et al. Rapid Prototyping of 3d DNA-Origami Shapes with Cadnano. *Nucleic Acids Research* **37**, 5001-5006 (2009).
3. Doty, D., Lee, B.L. & Stérin, T. in DNA 2020: Proceedings of the 26th International Meeting on DNA Computing and Molecular Programming, Vol. 174. (eds. C. Geary & M.J. Patitz) 9:1-9:17 (Schloss Dagstuhl--Leibniz-Zentrum für Informatik, 2020).
4. Wang, W. et al. Complex Wireframe DNA Nanostructures from Simple Building Blocks. *Nature Communications* **10**, 1067 (2019).
5. Fornace, M.E. et al. Nupack: Analysis and Design of Nucleic Acid Structures, Devices, and Systems. *ChemRxiv.* (2022).
6. Poppleton, E. et al. Oxdna: Coarse-Grained Simulations of Nucleic Acids Made Simple. *Journal of Open Source Software* **8**, 4693 (2023).
7. Poppleton, E., Romero, R., Mallya, A., Rovigatti, L. & Šulc, P. Oxdna.Org: A Public Webserver for Coarse-Grained Simulations of DNA and Rna Nanostructures. *Nucleic Acids Research* **49**, W491-W498 (2021).
8. Rovigatti, L., Šulc, P., Reguly, I.Z. & Romano, F. A Comparison between Parallelization Approaches in Molecular Dynamics Simulations on Gpus. *Journal of Computational Chemistry* **36**, 1-8 (2015).
9. Sample, M., Liu, H., Matthies, M. & Šulc, P. Hairygami: Analysis of DNA Nanostructures' Conformational Change Driven by Functionalizable Overhangs. *arXiv preprint arXiv:2302.09109* (2023).
10. Engel, M.C., Romano, F., Louis, A.A. & Doye, J.P.K. Measuring Internal Forces in Single-Stranded DNA: Application to a DNA Force Clamp. *Journal of Chemical Theory and Computation* **16**, 7764-7775 (2020).
11. Marko, J.F. & Cocco, S. The Micromechanics of DNA. *Physics World* **16**, 37 (2003).
